# Clonal diversification and histogenesis of malignant germ cell tumours

**DOI:** 10.1101/2022.02.01.478621

**Authors:** Thomas R. W. Oliver, Lia Chappell, Rashesh Sanghvi, Lauren Deighton, Naser Ansari-Pour, Stefan C. Dentro, Matthew D. Young, Tim H. H. Coorens, Hyunchul Jung, Tim Butler, Matthew D. C. Neville, Daniel Leongamornlert, Mathijs Sanders, Yvette Hooks, Alex Cagan, Thomas J. Mitchell, Isidro Cortes-Ciriano, Anne Y. Warren, David C. Wedge, Rakesh Heer, Nicholas Coleman, Matthew J. Murray, Peter J. Campbell, Raheleh Rahbari, Sam Behjati

**Author notes:** Joint first authors (Thomas R. W. Oliver, Lia Chappell).

## Abstract

Germ cell tumours (GCTs) are a collection of benign and malignant neoplasms derived from primordial germ cells. They are uniquely able to recapitulate embryonic and extraembryonic tissues, which carries prognostic and therapeutic significance. The developmental pathways underpinning GCT initiation and histogenesis are incompletely understood. Here, we studied the relationship of histogenesis and clonal diversification in GCTs by analysing the genome and transcriptome of 547 microdissected histological units. We found that the extensive diversification of tissues and genetic subclones were not correlated. However, we identified unifying features including the retention of fetal developmental transcripts across tissues, expression changes on chromosome 12p, and a conserved somatic evolutionary sequence of whole genome duplication followed by clonal diversification. Whilst this pattern was preserved across all GCTs, the developmental timing of the duplication varied between prepubertal and postpubertal cases. In addition, tumours of younger children exhibited distinct substitution signatures, including a novel one, which may lend themselves as potential biomarkers for risk stratification. Our findings portray the extensive diversification of GCT tissues and genetic subclones as randomly distributed, whilst identifying overarching tissue and tumour transcriptional and genomic features.

## INTRODUCTION

Germ cell tumours (GCTs) encompass a diverse spectrum of benign and malignant neoplasms of primordial germ cell (PGC) origin. Most malignant cases occur in the gonads of men aged 15-44 years old [1]. These tumours can be broadly divided by their histological composition into pure seminomas and nonseminomatous GCTs (NSGCTs). The latter, uniquely, can resemble a variety of tissues of embryonic and extraembryonic origin and are frequently composed of multiple histologies at once [2]. The histological composition of GCTs is of central significance for prognosis and patient management, with nonseminomatous GCTs conferring a worse prognosis [3, 4]. It is unknown whether histological diversification in an individual GCT is governed by regional genetic changes, as has been shown, for example, in different histological areas of the childhood kidney cancer, Wilms tumour [5]. Furthermore, the extent to which histogenesis in GCTs mirrors normal tissue development has not been established.

These questions may be answered by an investigation of GCTs that combines both genomic and transcriptional readouts. Previous efforts in this regard either utilised bulk tissues, where tissue-specific information within tumours containing mixed histological components would have been lost or studied microdissected tissue through limited assays such as targeted DNA sequencing [6–11]. The latter approach was guided by variants found in a single 40x whole genome taken from the primary tumour; an approach that likely lacks the necessary resolution to resolve tumour subclonality in detail [9]. Nevertheless, these efforts and others have yielded key insights about postpubertal testicular GCTs - irrespective of their broad histological category, they undergo whole genome duplication to form germ cell neoplasia *in situ* followed by the acquisition of further 12p copies to facilitate invasion [12].

Here, we examined the whole genomes and transcriptomes of 14 distinct histologies across 22 GCTs and four background normal testes at the resolution of individual histological units, to study the interplay of genetic diversification and tissue differentiation.

## RESULTS

### Overview of study

We assembled a primary cohort comprising 15 GCTs, encompassing 11 nonseminomatous GCTs, three pure seminomas and one case of a testicular prepubertal yolk sac tumour (median age 27 years, range 1 - 58) (**Extended data figure 1**). All samples were taken from the primary tumour. One case had only *in situ* disease available for analysis. Using laser capture microdissection, we excised 547 distinct histological units (DNA and RNA in 12/15 cases, DNA only in one case and RNA only in two; **Fig. 1a**, **Supplementary table 1**). This included 131 from tumours for DNA and 353 for RNA, as well as 63 microdissections from four regions of normal testis that served as a reference for the RNA experiments. In addition, we studied a further seven cases of prepubertal and peripubertal yolk sac tumours (age range 0.75 - 12 years, **Supplementary table 1**) by orthogonal bulk whole genome sequencing (WGS), to validate findings from the primary cohort.

**Figure 1.**
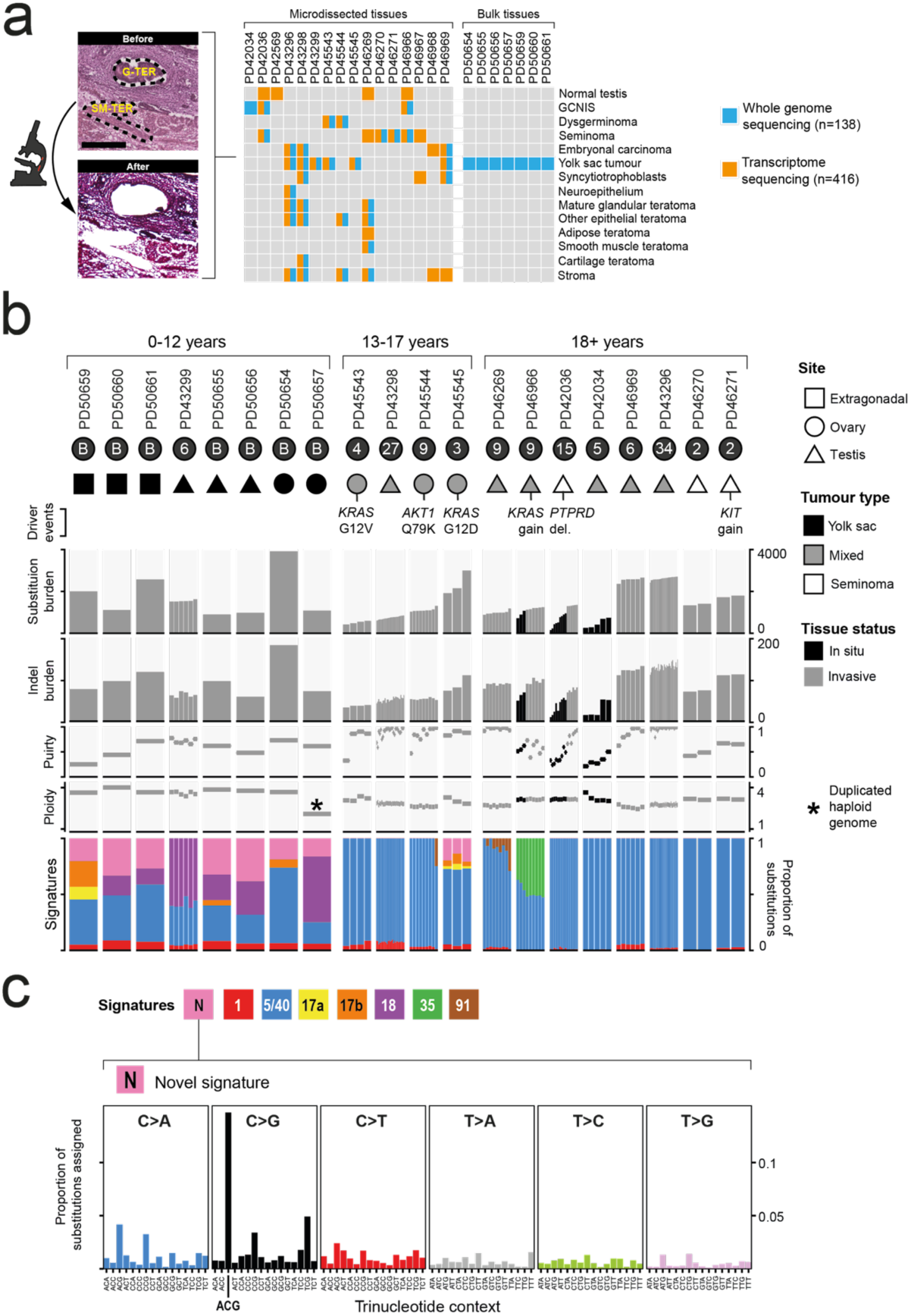
The mutational profile of GCTs. (**a**) Overview of the experimental design, including microphotographs from PD46269 illustrating different tumour histologies amenable to microdissection and sequencing. G-TER, glandular teratoma; SM-TER, smooth muscle teratoma. Scale bar represents 250 microns. The total number of whole genomes includes both the microdissected (131) and bulk (7) samples. Note that for the mixed tumours PD42034 and PD45545 only one histology was available for microdissection and PD42569 was normal testis from a healthy donor. (**b**) Summary plot of key genomic data and relevant metadata pertaining to each GCT analysed. Samples are ordered by patient and age. Each patient is labelled as having either a single bulk whole genome (B) or with the number of individual histological units microdissected. See Methods for driver annotation. (**c**) Trinucleotide context plot of the novel SBS signature detected.

### The somatic landscape of GCT microdissections

We generated WGS for 131 of the microdissected histological units, comprising 13 GCTs (**Fig. 1a**, **Supplementary table 1)**. The median coverage per tumour microdissection was 30 (range 15-48) (**Supplementary table 2**). All classes of somatic changes were called through an extensively validated variant calling pipeline **(Methods**, **Supplementary tables 2 - 6**). We calculated the median cancer cell fraction of substitutions called per dissection, corrected for copy number, to be 0.96 (range 0.81 - 1.08) (**Supplementary table 2, Extended data figure 2**), indicating that these microbiopsies largely represented monoclonal tumour cell clusters.

In terms of overall mutation burden and driver variants, the GCT microbiopsies broadly matched previous reports on postpubertal GCTs (**Fig. 1b**). In keeping with reports from exome analyses [6, 8], genomes of invasive GCTs exhibited 0.49 substitutions and 0.03 indels per Mb (**Supplementary table 2**). Within tumours that harboured both *in situ* and invasive disease, the former harboured fewer mutations (511 vs 1324 and 893 vs 1206 median substitutions for PD42036 and PD46966 respectively, p < 0.001 and p = 0.02, Wilcoxon rank sum test) (**Fig. 1b**). We identified typical GCT driver events in our cohort, including *AKT1* and *KRAS* substitutions, as well as gains of the *KRAS* and *KIT* oncogenes and a homozygous deletion of the tumour suppressor gene *PTPRD* (**Supplementary tables 4, 5, 7, 8**) [6, 8, 11]. Moreover, tumours exhibited typical GCT copy number profiles, as defined using 103 malignant testicular GCTs assembled by The Cancer Genome Atlas (**Methods**, **Extended data figure 3**) [6]. Taken together, these findings confirmed that our primary cohort largely represented classical GCT genomes, making our downstream analyses generalisable to GCTs more widely.

One exception to this was the case of a prepubertal yolk sac tumour (PD43299), which exhibited a single base substitution (SBS) signature not typically associated with GCTs. In postpubertal GCTs, most substitutions were, as expected, attributed to signatures representing errors of cell division (SBS5/40 and, to a lesser extent, SBS1) (**Methods**, **Extended data figure 4**, **Fig. 1b**, **Supplementary table 9**) or prior treatment (i.e., the platinum agent exposure SBS35 in case PD46966). By contrast, the prepubertal yolk sac tumour predominantly harboured substitutions assigned to SBS18. SBS18 is a non-ubiquitous signature that is most prevalent in normal human placental tissue and in the childhood cancers neuroblastoma and rhabdomyosarcoma [13, 14]. We pursued this finding through an orthogonal sequencing approach, using bulk tissues in an extended cohort of seven pre- and peripubertal testicular, ovarian and extragonadal yolk sac tumours. The results confirmed that pre- and peripubertal yolk sac tumours were distinguished by a contribution of SBS18. In addition, we discovered three further signatures heavily enriched in these cases; SBS17a, SBS17b and a novel signature characterised by C>G substitutions in an A[C>G]G trinucleotide context (signature “N” in **Fig. 1c**). Of note, the latter signature was absent from a recent meta-analysis of 23,829 cancer genomes [15]. The distinct profile of these cases was further underlined by the significantly higher burden of structural variants, including retrotransposition events, in patients aged 0-12 years vs those aged 18 or older (p < 0.001, Wilcoxon rank sum test) (**Extended data figure 5)**. Together, these genomic features indicated that mutational processes underpinning GCT formation may vary across age groups.

### Phylogeny and clonal diversification of tumours

To study the interplay between histogenesis and clonal diversification, we built phylogenetic trees for each microdissected tumour using somatic mutations (**Methods**, **Extended data figure 6**, **Supplementary table 10**). Our analyses were based on the principles that (i) mutations shared between different tumour regions define a common phylogenetic trunk and that (ii) copy number gains defining the trunk can be timed relative to the acquisition of point mutations. For example, mutations that occurred prior to duplication of an allele would be present on both derivatives, whereas variants generated after duplication would be confined to one copy only.

Overall, GCT formation across histologies and individuals was characterised by several recurrent features. Firstly, truncal mutations represented the majority of those found in each invasive tumour, suggesting subclonal diversification was a relatively late event and included all identified driver events (**Supplementary table 11**). Most copy number gains occurred within this trunk at a similar mutation time, consistent with whole genome duplication (**Methods, Extended data figure 7**). The duplication itself universally arose very early, with only ∼5.8 substitutions estimated to occur prior to this event across the entire genome of postpubertal (>12 years) GCTs (range 0 – 18) **(Methods,** **Fig. 2a, Supplementary table 12**). This makes postpubertal cases most unusual when compared with 2,096 cancer whole genomes across 31 tumour types analysed by the Pan-Cancer Analysis of Whole Genomes [16] (**Fig. 2b, Supplementary table 13**). In contrast, we estimated the median pre-duplication burden in patients aged 0-12 years to be 359.7 substitutions (range 10.6 - 788.8), suggesting that whole genome duplication, whilst a universal feature of GCT, occurs relatively late in the development in tumours of the young. No convincing evidence was found to indicate that the driver missense variants occurred prior to duplication (**Supplementary table 7**).

**Figure 2.**
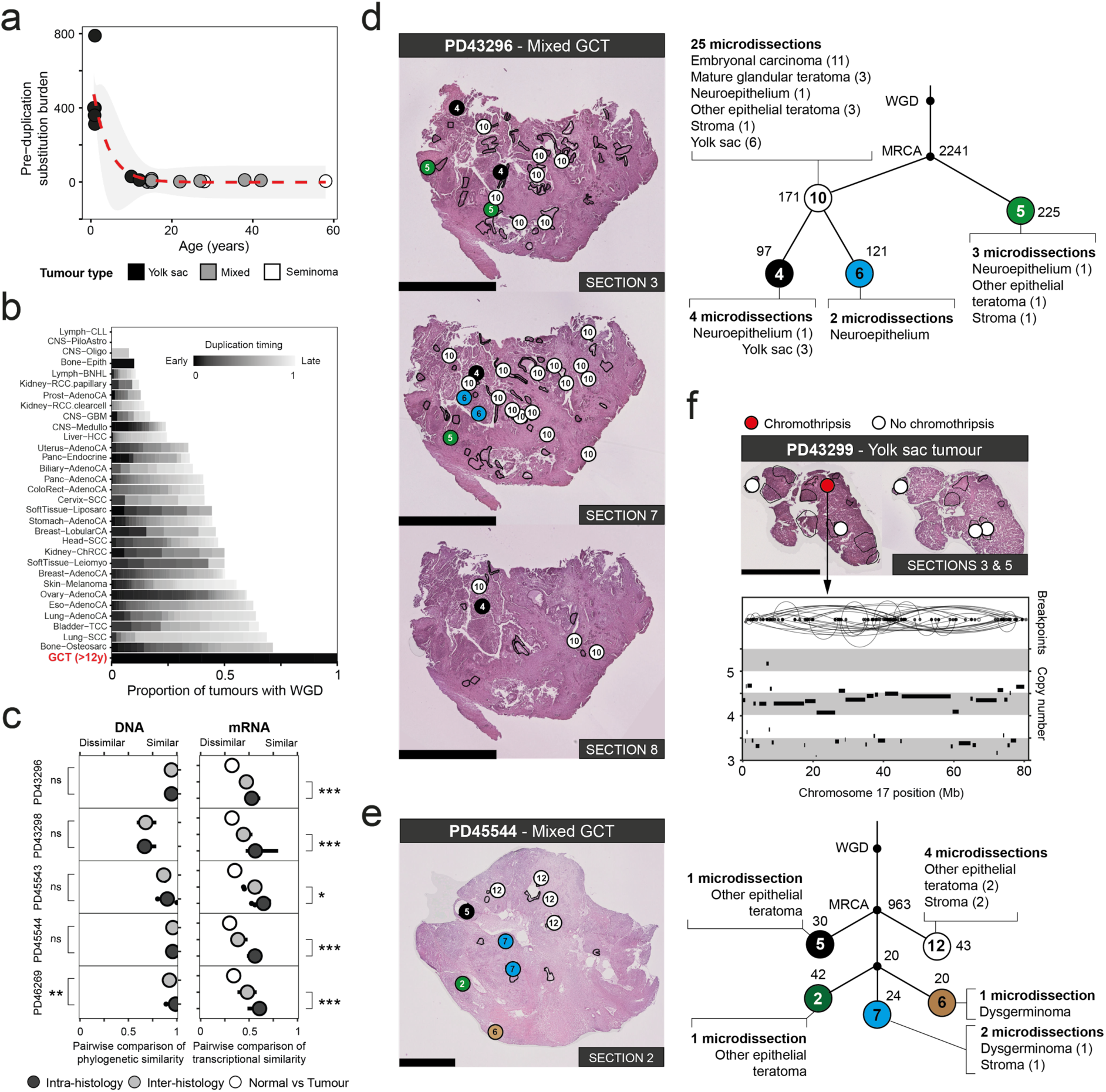
Whole genome duplication timing and tumour diversification. (**a**) Estimated burden of substitutions across the genome prior to the duplication event by age. The dashed red line is the fitted asymptotic regression with a surrounding grey ribbon indicating the 95% confidence intervals. The underlying equation is: pre-duplication substitution burden = -0.59 + (574.62 * e^(-e^-1.34 * age)). (**b**) Bar plot comparing the prevalence and timing of WGD between our postpubertal GCTs and the tumours analysed by PCAWG [16]. Tumour abbreviations used are as per the PCAWG studies. (**c**) Pairwise comparison of the genetic and transcriptomic similarity of microdissections within GCTs where multiple tissues underwent DNA and mRNA sequencing. Asterisks refer to p values derived from a permutation test using label-swapping, comparing each tumour’s intra- and inter-histology genomic and transcriptomic similarity. ns, not significant; *, < 0.05; **, < 0.01; ***, < 0.001. (**d - e**) Histological images from example testicular (**d**) and ovarian (**e**) mixed GCTs that underwent extensive multiregional sampling, each annotated with the mutation clusters that define the phylogenetic relationship of each microbiopsy. Circles on the histological images correspond to a numbered mutation cluster in the associated phylogeny on the right-hand side. The number next to each cluster (i.e., not within it) denotes the number of autosomal SNVs that support it. Circles are arbitrarily coloured to spatially highlight the clonal composition of each tumour. Each cluster is labelled with the number of microdissections for which it is the major clone and a list of the histologies the cluster pervades. MRCA, most recent common ancestor. These figures are simplified versions of the full phylogenies depicted in Extended data figure 6. (**f**) Subclonal chromothripsis of chromosome 17 in a prepubertal yolk sac tumour. Reconstructed breakpoints are illustrated above the copy number calls. All scale bars represent 2.5mm.

We next assessed when precisely during PGC development whole genome duplication occurred by dividing the pre-duplication substitution burden estimates by the reported mutation rate per cell division within PGCs [17]. We estimated the median postpubertal GCT whole genome duplication occurred at ∼10 cell divisions post-PGC specification (range 0-36, lower and upper bounds of mutations per cell division estimate) (**Methods**), overall placing the genetic hallmark of GCT initiation - whole genome duplication - in fetal life for many tumours.

### Relation of histogenesis to clonal diversification

Next, we asked whether histogenesis and genetic diversification were correlated by analysing both DNA and RNA from the same histological units. We generated cDNA libraries through a modified single cell mRNA sequencing protocol (**Methods**). 112 out of 131 whole genomes derived from microdissections had transcriptomic data from adjacent regions available. These libraries yielded expression profiles for 55,502 non-mitochondrial features with a median 298,080 reads mapped per microbiopsy (range 13,822 - 3,863,511 reads) (**Supplementary table 14**). We performed pairwise comparisons to identify the similarities of genomes and transcriptional profiles within each GCT (**Methods,** **Fig. 2c**). We considered comparisons of the same and of different histologies within each tumour. The results suggested that there was no difference in the genetic similarity between biopsies of the same or different histology in most cases. Instead, genomic diversification followed an anatomical pattern where, within each patch of a unique somatic subclone, various histologies had been generated, as our reconstructed phylogenies elucidated (**Fig. 2d - e, Extended data figure 6, Supplementary table 10**). By contrast, the transcriptional expression within a single GCT histology was significantly more similar compared with other tissues, indicating that histology-specific, protein-coding transcription transcended genetic heterogeneity (**Fig. 2c**). A striking example was seen in a pure yolk sac tumour in which one subclone was defined by 66 structural variants rearranging chromosome 17 (chromothripsis) which generated potential driver events (e.g., loss of *TP53*), as previously described in an osteosarcoma (**Fig. 2f**) [18]. However, chromothripsis did not significantly perturb the global transcriptional proximity between different tumour clones (p > 0.05, permutation test) (**Methods**, **Extended data figure 8**). In aggregate, our findings indicate that histogenesis in GCTs is not governed by somatic genetic diversification. This finding corroborates a previous observation in bulk data of transcriptional clustering by gross histological category despite the absence of an apparent unifying genomic event [6].

### Fetal signals underpin GCT histogenesis

A fundamental question of GCT histogenesis is whether differentiated tissues, such as a region of GCT cartilage, show abnormal gene expression relative to their adult, normal counterparts. We addressed this question by comparing transcriptomes of GCT tissues with reference transcriptomes of corresponding fetal and mature tissues, as defined by single cell mRNA sequencing. We examined a total of 416 histological units from 14 tumours and four regions of histologically normal testes covering a total of 14 histological structures (**Fig. 1a**, **Extended data figure 1**, **Supplementary table 1**).

Using single cell reference data from fetal and adult tissues corresponding to each GCT tissue [19–23], we found that tumours not only expressed lineage-specific transcripts but also consistently retained expression of fetus-specific features (**Fig. 3a - c**). This remained the case even in apparently mature tissues such as the smooth muscle teratoma where *IGF2*, typically restricted to high expression in the fetus [24], was readily detectable alongside the typical smooth muscle markers *ACTA2*, *MYH11* and *TAGLN* [23]. In contrast, the microdissected adult seminiferous tubules did not demonstrate the fetal signal seen in the malignant tissues recapitulating primordial germ cells, i.e. seminoma, dysgerminoma and GCNIS (**Fig. 3a**).

**Figure 3.**
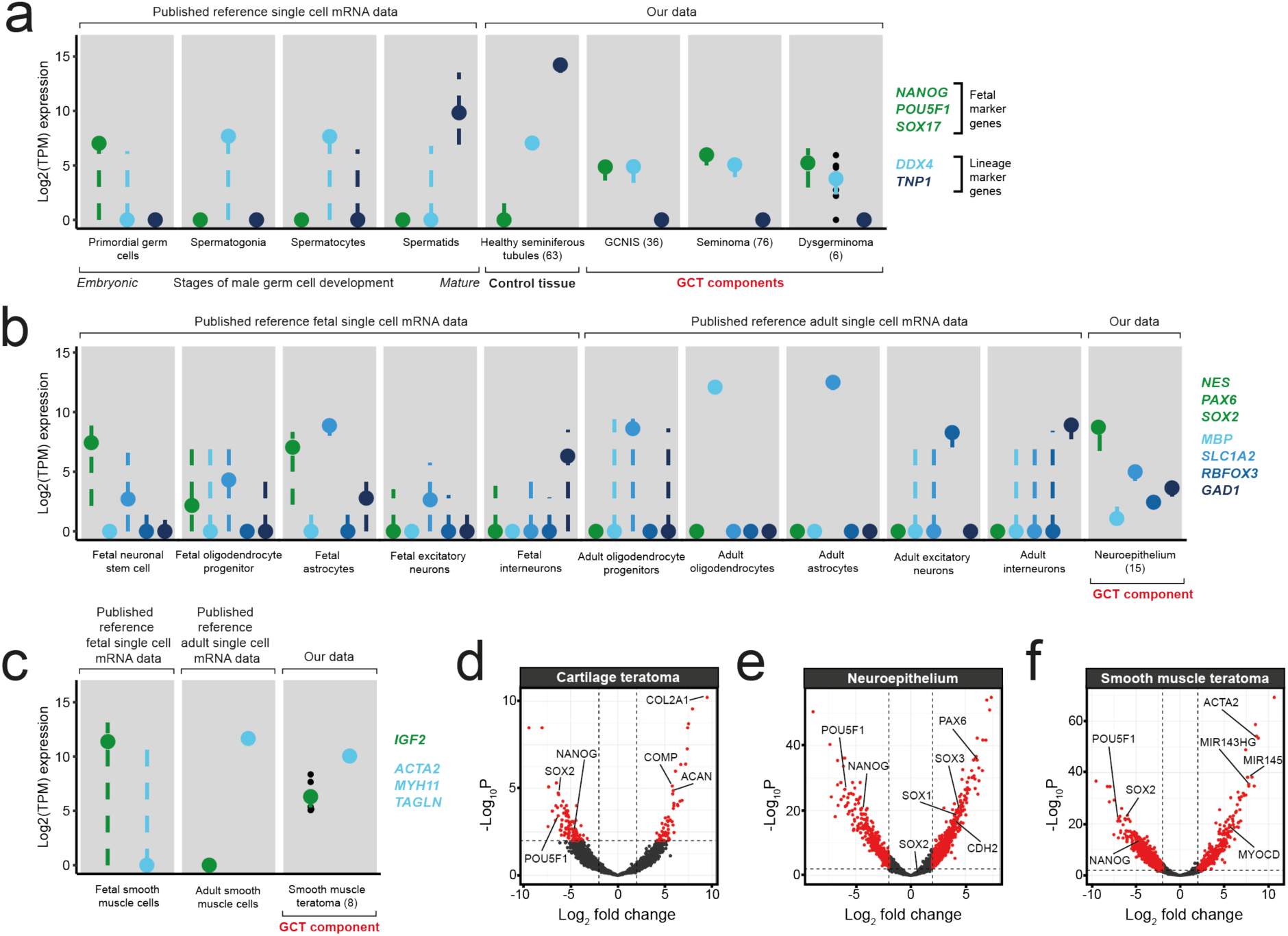
Pathways of GCT histogenesis. (**a - c**) Comparison of the expression of marker genes in example GCT tissues and their corresponding fetal and adult counterparts, using reference single cell datasets [19–23]. Coloured dots denote the median and dashed lines the interquartile range. Individual data points are plotted where the total data points supporting a point range are ≤10. Each point represents the expression of a single gene per tissue. (**d - f**) Example volcano plots illustrating the differential expression analysis between embryonal carcinoma and more mature GCT tissues. The dashed lines represent the cut-offs for the log2 fold change (> |2|) and adjusted p value (<0.01, Benjamini-Hochberg correction) considered significant. Genes enriched in each differentiated tissue are shifted to the right. Marker genes used to define embryonic stem cells and each differentiated tissue are annotated on plot. The full list of genes and tissues comparisons can be found in Supplementary table 15.

If the transcriptomes of GCT tissues resembled their fetal correlates, we might then better appreciate this by performing differential expression analysis between each individual NSGCT component and the NSGCT subtype embryonal carcinoma, which phenotypically recapitulates human embryonic stem cells (hESCs). (**Methods, Supplementary table 15**) [25, 26]. Here, illustrated through use of hESCs markers, as well as those corresponding to each differentiated tissue, we found gradients of relative expression akin to those seen in embryogenesis (**Fig. 3d - f**). Maturing tumour tissues overexpressed genes known to regulate cell fate specification to their matching fetal tissues, such as in neuroepithelium (*SOX1*, *SOX3* and *PAX6*) and smooth muscle (*MYOCD*, *MIR143HG* and *MIR145*) [27–30]. This lineage specification is likely to be determined at an epigenetic level, as has been suggested previously [31].

### A canonical GCT transcriptome

Despite the histological and protein-coding transcriptional diversity of GCTs, it is conceivable that there is a global component of the transcriptome that transcends tissues and tumours. We found significant enrichment of chromosome 12p gene expression across the invasive tumour histologies (**Fig. 4a**, **Supplementary table 16**), by assessing the expression profile of each GCT tissue, relative to normal testis within each cytoband along the genome (**Methods**). The enrichment corresponded to copy number gain, with 12p genes generally possessing a significantly higher log2 fold-change relative to healthy seminiferous tubules than regions nearer baseline ploidy (p < 10^-5^, permutation test) (**Methods**, **Fig. 4b**, **Extended data figure 9**). This finding was of particular importance and plausibility as gains of chromosome 12p (usually arranged in an isochromosome) are a near universal feature of postpubertal GCTs and is consistent with previous bulk sequencing of GCTs and other malignancies [32, 33]. We note that the mapping of the transcriptional change to copy number was not exact and may be explained in part by use of relative expression to normal testis and the influence of gene promoters that lie outside of the region gained. Examining the expression of individual genes along 12p revealed the *bona fide* oncogene *CCND2* to be universally overexpressed in all invasive tumour histologies investigated using our conservative cut-offs (≥2 log2 fold-change and <0.01 adjusted p value), as well as *ATN1* and *PTMS*. *KRAS*, in contrast, was only overexpressed in embryonal carcinoma (**Supplementary table 17**). PTMS (parathymosin) is thought to facilitate chromatin remodelling through interaction with the linker histone H1 and can induce human sperm nuclei to undergo decondensation, possibly implicating it in GCT epigenetic remodelling [34]. Other features that were widely expressed (*SLC2A3*, *PHB2*) were consistent with previous reports [32]. Together, a picture emerged confirming a conserved GCT transcriptome in the form of 12p gene overexpression, regardless of invasive histology, driven by the defining 12p gain.

**Figure 4.**
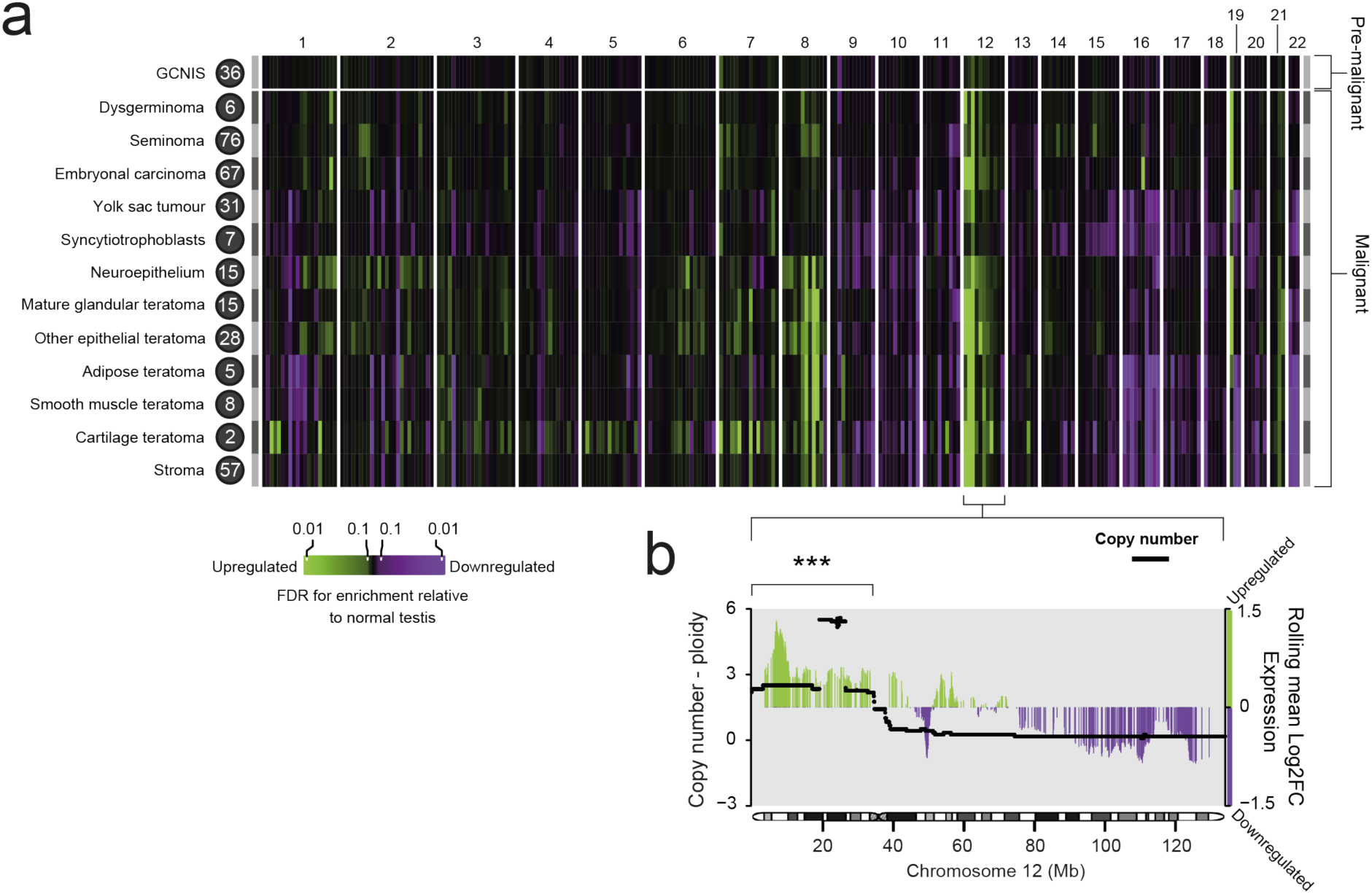
The relationship between GCT genome and transcriptome. (**a**) Heatmap showing gene enrichment per GCT tissue relative to healthy seminiferous tubules, binned by cytoband. Colours correspond to significance of enrichment according to the adjusted p value (false discovery rate or Benjamini-Hochberg correction). The number next to each histology is the number of eligible microbiopsies that informs the analysis. (**b**) Combined plot of the chromosome 12 copy number changes across all invasive tumours and the rolling average log2 fold change in gene expression compared with healthy seminiferous tubules. Window size for the rolling average is 50 genes. The average log2 fold change in expression across 12p was significantly higher than across comparable numbers of genes found across regions with near baseline ploidy (p < 1 x 10^-5^) (**Extended data figure 9**), as indicated by the asterisks.

## DISCUSSION

We studied the origins and tissue diversification of GCTs from DNA and mRNA sequences derived from microdissected histological units. Our approach enabled us to directly overlay anatomical boundaries of clonal diversification with histological features. Our result revealed that histogenesis in GCTs is not demarcated by clonal territories. Instead, GCT tissues appear to arise independently of somatic diversification along transcriptional pathways that broadly mirror normal human tissue development. An important distinction from normal histogenesis was the retention of fetal developmental genes in GCT tissues, even in seemingly mature tissues, that emerged as a defining feature of GCT components. It is conceivable that this incomplete maturation represents a targetable vulnerability for interventions that promote differentiation, akin to therapeutic efforts in childhood cancer that aim to overcome maturation blocks [35–38]. Instances of more aggressive, differentiated GCTs are rare, such as non-gestational choriocarcinoma which may reflect the normal invasive potential of the trophoblasts they recapitulate [39]. Beyond the retention of fetal transcripts, we identified two further stereotypical features of GCT tissues. First, we observed dysregulation of gene expression on chromosome 12p, amplification of which is a key genomic hallmark of GCT, to be conserved across postpubertal GCT tissues. Second, the somatic genetic development of GCT genomes exhibited extensive diversification yet was unified by a common root that lay in WGD.

It has previously been suggested that pre- and postpubertal GCTs may arise from different stages of PGC specification [40]. The key evidence supporting this notion includes variations in the tissue composition of GCTs across different age groups, a reduced prevalence of 12p gain in tumours of the young, and nuances in translational protein-coding gene profiles [41–44]. Our investigation has revealed stark somatic genetic differences between pre- and postpubertal GCTs. Whilst the near universal step of WGD was preserved across all GCTs, it consistently occurred later in tumours of the young although the exact mechanisms by which it occurs in each case requires further investigation. Furthermore, our analyses revealed distinct mutational signatures that underpin many substitutions in GCTs of young children. These findings would support the view of a fundamental developmental distinction between pre- and postpubertal GCTs that may delineate a subgroup of ‘true’ paediatric tumours which may be of clinical relevance. A major challenge in the clinical management of GCTs remains whether to treat older, peripubertal children according to ‘paediatric’, less intense treatment protocols, or as ‘adults’ who generally require more intensive cytotoxic chemotherapy [45]. We speculate that timing of whole genome duplication and distinct mutational signature profiles may lend themselves as potential tangible biomarkers for treatment stratification in this context.

Our investigation has directly addressed the relationship between genetic and transcriptional diversity in GCTs. Although limited by the relatively few cases that possessed each single histology or many simultaneously, the high-resolution multi-omics approach of our study seems to indicate that the superficially heterogeneous GCTs still possessed canonical transcriptional and genetic features that underpin GCT development. We would expect our approach to be broadly applicable across human cancer to study the interplay of somatic changes and transcription at the level of microscopic tumour regions.

## METHODS

### Tissue collection, handling and microdissection

Samples were stored fresh frozen after collection in accordance with protocols approved by the relevant local ethics committee (UK REC approval reference numbers 03/018, 08/h0405/22+5, 12/NE/0395, 16/EE/0394, 17/LO/1801,18/EM/0134 and 18/NW/0092). The tissues in the primary cohort were fixed using PAXgene fixative (Qiagen) and prepared for microdissection as described previously [46]. To ensure that each component of the tumour isolated was correctly identified, a reference H&E slide and slides stained with the relevant immunohistochemical antibodies were generated and reviewed by a consultant histopathologist (**Extended data figure 1**).

Each sample was taken from the primary tumour. The age at diagnosis was provided for each patient.

### DNA library preparation and sequencing

Low-input, whole genome sequencing libraries were created from each microbiopsy following an established protocol [46]. 150bp paired-end reads were then generated from these libraries using either the Illumina HiSeq XTEN or NovaSeq sequencing platforms to a target coverage of 30x. For the additional bulk samples, all seven were sequenced on the NovaSeq platform only (**Supplementary table 2**). These reads were aligned to the GRCh37d5 human reference genome using the Burrows-Wheeler Aligner (BWA-MEM) [47].

### RNA library preparation and sequencing

Microbiopsies were collected in wells pre-filled with 50µl of RLT Plus lysis buffer (Qiagen) during microdissection. Regions where microdissections had been taken for DNA sequencing were specifically targeted, using the adjacent tissue sections. RNA microdissections were considered “adjacent” if they were <1mm from where a successful DNA library had been generated and were of the same histological feature (**Supplementary table 2**).

AMpure XP beads (Beckman Coulter) were used to recover RNA from the lysate. Oligos and reagents from the Smart-seq2 protocol [48] were used to reverse transcribe and amplify full-length cDNA. Libraries were prepared from the cDNA using the NEBNext Ultra II FS DNA Library Prep Kit for Illumina (New England Biolabs). Libraries were sequenced with the Illumina HiSeq 2500 or 4000 systems to produce 100bp paired-end reads. Adaptors and low quality reads were removed using Trim Galore (https://github.com/FelixKrueger/TrimGalore) and following parameters: -q 20 --fastqc --paired --stringency 1 --length 20 -e 0.1 The Spliced Transcripts Alignment to a Reference (STAR) aligner was used to map the raw sequencing reads to the GRCh37d5 human reference genome [49].

### Estimating contamination of tumour samples

To exclude contamination as a source of false positive variant calls, we ran the Conpair algorithm [50] on all whole genome data included.

### Estimating the callable length of the tumour genome

When calculating the mutation burden per megabase, we adjusted for the length of the genome we were able to call mutations across. We ran Mosdepth [51] to exclude loci that have fewer than 4 reads covering them and then removed regions masked by the variant callers due to their highly repetitive nature.

### Variant calling and filtering

All DNA mutation calling was performed against a matched normal blood or non-neoplastic testis microbiopsy sample.

Substitutions were called using the CaVEMan algorithm [52]. We only kept candidate variants if the reads supporting them passed a minimum median alignment score (≥140) and fewer than half were clipped. For bulk samples, variants needed to be supported by ≥4 variant reads with a total coverage of ≥10x. Further artefacts resulting from the low input DNA library preparation method in the microdissected samples were then removed using previously validated filters [46]. Low input pipeline substitution artefacts are usually found on reads that share a similar alignment. Consequently, these filters are based on statistics such as a variant’s position compared to the start of alignment and its standard deviation and median absolute deviation on the reads supporting it. To improve our sensitivity for calling low depth variants in any given microbiopsy, a particular problem for the lower purity and aneuploid *in situ* disease, we leveraged the multi-sampling nature of the experiment by conducting a pileup of all substitutions called across all patient samples with minimum base (25) and read mapping (30) quality thresholds. Putative mutations found at loci with universally low coverage (mean ≤10x) across a patient’s samples were then removed as probable mapping artefacts. Finally, to differentiate true low depth variants from sequencing artefact, we fitted a beta-binomial model adapted from the Shearwater variant caller [53] to establish a locus-specific error rate using a reference panel of 250 unrelated, normal samples that have been subject to the same library preparation and sequencing platforms [13, 54, 55]. A variant was considered real in a given sample if p < 0.001 after multiple-test correction (Benjamini-Hochberg method). To ensure the veracity of our variant calling from this pipeline, a subset of substitutions were visually inspected using the genome browser Jbrowse [56].

Small indels were identified using the Pindel algorithm [57]. All putative indels had to pass a minimum quality score (≥300), not fall within 1bp of a SNP or 5bp of an indel call found in the matched normal sample and not be a 1bp indel at a homopolymer run of 7bp or longer. Indels were then only considered if ≥5 variant reads supported them. For bulk samples, a total coverage at the indel site of ≥10x was required, as well as a minimum of at least one read supporting it in both directions. Indels from microbiopsies underwent similar genotyping to that described for substitutions. Genotyped variants were then filtered to remove those with a low average coverage (mean ≤10x) before being subjected to the aforementioned Shearwater- like approach. Due to computational constraints, the unmatched normal reference panel was reduced to 100 samples for indel filtering. Tumour variants seen at a VAF of ≥0.2 in the matched normal or across this unmatched panel were filtered out. Mutations were excluded if the combined sum of unknown and ambiguous reads at that position across a patient’s samples was greater than the number of variant reads or if half or more total reads at that locus were of low quality (mapping quality <20), as previously described [54, 56]. Once more, inspection of a subset of the final indel calls in Jbrowse [56] validated this stringent filtering approach.

Battenberg [58] was used to call copy number aberrations and estimate the purity and ploidy. As Battenberg is primarily informed by the B allele frequency when determining the copy number state, very high purity tumour samples - such as those obtained by microdissection - with regions of loss of heterozygosity can be challenging to accurately call. To overcome this, Battenberg was run twice - once as a standard tumour and matched normal pair and the second time merging the tumour and normal sample BAM files to “lower” the tumour purity. The results from both runs were manually reviewed and subject to the DPClust algorithm [58] which determines the subclonal architecture of a sample using both the substitutions and copy number calls. The best fit according to this combined approach was kept. Where the merged BAM file Battenberg calls were kept, the purity from the original run was still used in downstream analyses. The DPClust algorithm was run for 10,000 iterations with the first 3,000 dropped as burn-in.

Structural variants were detected using the BRASS algorithm [58]. False positive calls because of the low input DNA library preparation were removed using a previously described approach [59] which was informed by the same 100-sample normal reference panel used in indel filtering. Further artefacts were flagged by manually reviewing the borderline calls that narrowly passed filtering and removing them. Additional filtering for the bulk samples consisted of removing any variants also called in the matched normal. The final step for both cohorts was to remove any remaining variants found within 100bp of any retrotransposition event.

We called somatic retrotransposition events using a separate pipeline to the one detailed above. First, those with exact breakpoints were identified using the TraFic algorithm [60]. To then identify further events without breakpoints, the raw calls from the BRASS algorithm were examined to identify read clusters associated with known L1 germline sources. All putative variants were then manually reviewed. The final calls can be found in Supplementary table 6.

### Mutational signature extraction

*De novo* single base substitution (SBS) signatures were initially extracted using the hierarchical Dirichlet process (HDP version 0.1.5, https://github.com/nicolaroberts/hdp) [61]. This was done on a per patient basis, i.e. all unique substitutions across a tumour, to prevent double counting. This was run across 20 independent posterior sampling chains with 80,000 burn-ins and another 20,000 sampled iterations. The resultant signatures were deconvolved against the COSMIC version 3.2 SBS signatures using a previously described expectation-maximisation algorithm [62]. The novel signature detected in the initial run showed clear conflation with SBS1, likely as a consequence of higher SBS1 burdens and the novel signature colocalising to the same samples (**Extended data figure 10**). To clean this up, HDP was run once more using only the prior COSMIC signatures, including SBS1, found in the initial run. The resultant novel signature is shown in Figure 1c. All the components extracted at the final step can be seen in Extended data figure 10. Signatures were then mapped back to individual samples using SigProfiler (version 1.015) [15]. SigProfiler performed an independent *de novo* signature extraction whose output was decomposed using the final list of HDP signatures. For the SigProfiler *de novo* extraction step, sigProfilerExtractor was run using a minimum and maximum of 2 and 15 signatures respectively with all other parameters set as default.

### Driver mutation analysis

Driver mutations were initially considered in known cancer genes, as defined by the COSMIC version 94 cancer genes consensus. Missense mutations and in frame indels that occurred within genes annotated as oncogenes, or who act in a dominant manner, and were found at previously documented hotspots were considered driver events. For recessive cancer genes, all intact gene copies had to be lost to consider it a driver event. This included substitutions, small indels, structural variant breakpoints that truncated the gene footprint and copy number changes. We defined focal amplification and deletion driver events according to previously outlined heuristics [58]. Briefly, gained segments had to be <1Mb in size and a minimum of five or nine intact copies of the oncogene had to be gained, depending on whether the average ploidy was below or above 2.7. Focal deletions shared the same segment size and ploidy cut- offs but with a total copy number of zero or less than average ploidy minus 2.7.

The *KRAS* and *KIT* gains noted in Figure 1b did not fulfil this definition for focal amplification, hence their notation as “gain”, rather than “amplification”. Both possessed the minimum number of copies, however they lay on segments between 1 and 10Mb in size. They were retained and noted, however, in view of their role as classic GCT drivers.

As a second, systematic check for drivers, we employed two computational methods. The first, dNdS, tested for genes under positive selection [63]. This did not yield any significantly enriched genes, likely due to the modest cohort size. Furthermore, the CHASMplus algorithm [64] was then run to systematically identify driver substitutions across the cohort with the three previously identified *KRAS* and *AKT1* mutations scoring most highly (**Supplementary table 8**).

### Aggregated copy number comparison against TCGA data

Using only GCT microbiopsies from invasive tissues, i.e. not GCNIS, with a purity >40%, the major clone copy number state (as defined by Battenberg) was extracted per 10kb genome bin. The purity threshold was used to ensure confidence in the copy number calls. The median total copy number across all microbiopsies was used to generate per tumour copy number profiles from which the median tumour ploidy was subtracted to distinguish additional gains for their higher starting copy number state baseline because of WGD. The average of this value was used in a cohort-level aggregated copy number profile. The prepubertal case (PD43299) was excluded from this analysis in view of its higher overall ploidy and lack of comparable samples in TCGA reference [6].

For the comparison with TCGA data [6], we only included TCGA samples with a purity greater than 40%, leaving 103 eligible tumours.

### Detection of chromothripsis

Chromothripsis-like events were detected using ShatterSheek [65]. Putative calls, both high and low confidence, were manually reviewed to remove false positive calls.

### Tumour phylogeny reconstruction

Tree building was performed using only samples with an average reads per chromosome copy of ≥5. The average reads per chromosome copy in a given sample was calculated using the following formula [16]:

Average_reads_per_chromosome_copy = (purity) / ((purity * ploidy) + ((1 - purity) * 2)) * (tumour_coverage)

The approach to phylogenetic tree reconstruction depended upon the number of samples available from a given tumour. For five samples or fewer, we used the DPClust algorithm [58] with the Gibbs sampler run for 3,000 iterations and the first 1,000 dropped as burn-in. More highly sampled tumours were subject to multidimensional DPClust instead [66]. This iteratively compares the phylogenetic distance of sample triplets before combining all outputs into a single unifying tree structure. Due to computational constraints the iterations and burn-ins used by the Gibbs sampler here were adjusted to 1,000 and 200 respectively. Both methods were informed by both the substitutions and copy number aberrations. Clusters accounting for <1% of total substitutions, fewer than 20 substitutions or small clusters that violated the remainder of the tree were removed (e.g. residual sequencing artefact that was found at low VAF across all mutation clusters) (**Supplementary table 10**). Only autosomal substitutions were included in this analysis.

### Estimating the proportion of substitutions within the tumour’s phylogenetic trunk

The tumour phylogenies generated served as the input to our calculation of the proportion of substitutions found within the phylogenetic trunk. This approach had the advantage of accounting for substitutions that may be apparently absent in a given sample but were in fact lost through a subsequent deletion or chromosomal loss. The proportion was calculated using the following formula:

Proportion_subs_in_trunk = (n_subs_in_trunk) / ((n_subs_in_trunk) + (sum(subclonal_branch_length) / (n_subclonal_branches)))

For tumours with GCNIS, the mutations shared between GCNIS and the invasive tumour were included in the trunk. To calculate the average branch length from the trunk, the summed mutation distance between the trunk and each branch tip was divided by the total number of branch tips. This meant double-counting some mutations in cases where two or more branch tips shared mutations not found within the phylogenetic trunk to normalise the lengths of all subclonal branches emanating from the trunk equally (**Supplementary table 11**).

### Timing copy number events

To ensure each copy of a chromosome within a sample had sufficient coverage to accurately time clonal copy number changes along it, we limited our analysis to the 96 microbiopsies and 6 bulk samples with a calculated average of ≥7 reads per chromosome copy. We then ran mutationtimeR [16] to identify clonal substitutions and estimate the probability that they each occurred prior to a copy gain. All analysed samples were determined to have undergone whole genome duplication as their ploidy (weighted by subclonality) was ≥2.9 - 2 * (Extent of homozygosity, weighted by subclonality), the approach taken by the PCAWG Consortium [16].

To estimate the substitution burden prior to WGD, we first identified all 2+0, 2+1 and 2+2 copy number segments that were predicted to be involved in WGD. We excluded low confidence segments, defined arbitrarily as those with ≥0.5 width confidence intervals for the mutation time estimate of their gain. All substitutions predicted to occur prior to duplication across these segments were then added together and adjusted for the copy number configuration of the tumour:

CN_adjusted_burden = sum(pre_dup_subs) + (sum(pre_dup_subs) * (sum(total_subs_2+1,2+0_) / (sum(total_subs_2+2,2+1,2+0_))

Pre_dup_subs represents our raw pre-duplication substitution counts whilst total_subs is all substitutions called across the eligible segments matching the configurations listed in the subscript. This adjustment accounts for the pre-duplication substitutions on the minor allele in regions of 2+0 or 2+1 that cannot be identified. Lastly, we extrapolated this adjusted value to a genome-wide estimate, according to how many bases were covered by our included copy number segments.

To convert the substitution burden to a number of cell divisions, we used a previously published estimate of 0.5-0.7 substitutions incurred per cell division within PGCs [17]. We anticipated that, by running our variant calling against a matched normal sample, that the earliest embryonic mutations (typically 1 - 2) [13, 67, 68] would not be detectable by our pipeline and thus most detectable mutations would have emerged post-PGC specification.

We considered alternative timing methods too, such as placing WGD in chronological time by restricting analyses to C>T substitutions in a CpG dinucleotide context, as was done in PCAWG [16]. The reasoning here was that these mutations are thought to accrue in a linear, clock-like fashion whilst other mutational processes will fluctuate over the course of a tumour’s life, making real-time estimations of copy number gain events from mutation time more difficult. We deemed this approach unsuitable for our study cohort, however. GCTs are characterised by low tumour mutation burden, meaning that restricting analyses to such a limited mutation context would leave very few mutations to analyse. Perhaps more importantly though, GCNIS and seminoma are characterised by global hypomethylation which may be re-established in NSGCTs [6, 69]. This is likely to influence the mutation rate at CpG sites considerably as mutations in this context are usually the consequence of spontaneous deamination of 5-methylcytosine, typically causing SBS1-pattern mutations [15]. With such dynamic changes in the methylation of the GCT genome, it cannot be assumed that the mutation rate at these sites in GCTs is linear.

Only tumour types with 10 or more samples were considered within the PCAWG dataset [16] when comparing it to our postpubertal GCT cohort.

### RNA sequencing data pre-processing

500 microdissections were initially taken from across the 14 tumours and 4 regions of histologically normal testis. 40 were excluded due to contamination by other tissues during microdissection, ambiguity over their histological categorisation, or because they were neither neoplastic nor seminiferous tubules (e.g. populations of Leydig cells or lymphocytic infiltrates). The library depths achieved across the 55,502 initially mapped nuclear features were somewhere between those seen in single cell and standard bulk RNA experiments with a median of 246,980 mapped reads (range 682 - 3,863,511, interquartile range 86,612 - 651,605) (**Extended data figure 11**). Reads were counted per feature using featureCounts (version 1.5.1) [70].

To filter out failed or low-quality libraries, we considered genes to be expressed if at least five reads were mapped to them and assessed how many genes were expressed per microbiopsy (**Extended data figure 12**). The median number of features expressed to this depth per sample was 8,916 (range 0-22,865, interquartile range 4,150-13,654). After removing 44 microbiopsies which expressed fewer than 1,000 features to this depth, 416 were left for downstream analyses.

To assess the mRNA quality of the remaining samples, we inspected a uniform manifold approximation and projection (UMAP) of their expression using the Seurat package [71] (**Extended data figure 13**). Histology was a strong determinant of sample clustering although we found embryonal carcinoma in particular retained patient-specific signals. We found no evidence of batch effect. We used marker genes from previously described GCT histologies to further assess the data quality and found their expression to localise to the expected tissues [72–75] (**Extended data figure 14**).

### Differential expression analyses

The Limma-Voom method [76] was used to perform all differential expression analyses. The default parameters were used to filter genes by a minimum expression within a test group necessary for differential expression to be confidently determined.

To identify the significantly enriched and depleted regions of gene expression across each GCT histology isolated, differential expression analysis was performed between each tissue and the healthy seminiferous tubule reference microbiopsies. Changes in expression at the level of the cytoband could then be confirmed by gene enrichment analysis using Limma’s ‘camera’ function and the MSigDB C1 gene set [77].

A pan-GCT expression profile for chromosome 12 was constructed by performing differential expression analysis on all invasive GCT tissues together against healthy seminiferous tubules, with the histological subtype set as the blocking factor (random effect) to adjust for histology-specific variability in 12p gene expression. In order to establish whether the high expression across 12p - which was universally gained in tumours that had undergone both DNA and RNA sequencing - could be due to chance, we performed 100,000 random samples of genes from genomic regions near baseline ploidy and measured their average log2 fold change in expression and compared them to the average across 12p. ‘Near baseline’ was arbitrarily set as ±0.5 away from the Battenberg ploidy estimate. Each sample contained 226 genes, reflecting the 226 genes that lay on 12p which were retained during the pan-GCT differential expression analysis. The remainder were filtered out due to a paucity of mapped reads in both the normal testis and GCT samples.

### Pairwise genetic and transcriptomic similarity scores

For tumours where DNA and mRNA sequencing for at least 2 invasive histologies were available we devised scores to indicate how similar a given pair of microdissections were. The genetic score was derived from the proportion of substitutions that the two cuts share, compared to their average burden, as determined by the mutation clusters called across each tumour (see “Tumour phylogeny reconstruction”). A mutation cluster was considered present in a sample if its cancer cell fraction was > 0.1.

For the transcriptomic similarity, we compared only protein-coding genes and excluded gene sets that were not of interest or likely to add noise to the analysis, including haemoglobin, immunoglobulin, cycling and housekeeping genes. Subsequently, any given two samples were compared by measuring the Pearson correlation coefficient between them. PD46969, although possessing both DNA and mRNA for multiple invasive GCT histologies, was excluded from this analysis on the basis that multiple samples were available for microdissection but the tissues isolated were not well represented across all biopsies. For example, yolk sac tumour DNA sequences were all derived from one biopsy and syncytiotrophoblasts from the other. Such large spatial biases were likely to confound assessment of the relationship between histology and phylogeny.

To assess the significance of the differences in genomic and transcriptomic similarity we observe between the intra- and inter-histological scores, we randomly swapped the labels for all pairwise comparisons in a tumour 1,000 times and plotted the difference in the median scores generated. This permutation approach provided a distribution from which a p value could be derived. For PD45543, which only had four whole genomes, 15 combinations would have exhausted the random sampling space, so we compared the observed genomic difference to the 14 possible alternatives.

### Global transcriptional effects of chromothripsis

To assess whether subclonal chromothripsis of chromosome 17 in PD43299, a pure yolk sac tumour, had a significant impact on the transcriptome, we examined the intra-tumoral Pearson correlation between regions of yolk sac tumour. 6 tumours were included. Our hypothesis was that chromothripsis would increase transcriptional heterogeneity which could be measured through an increase in variance. After calculating the variance of yolk sac tumour transcriptional similarity per individual, 1000 iterations of random sampling of 6 correlations, matching the size of the PD43299 correlation matrix, generated a distribution of variance against which we could compare the variance of transcriptional similarity in PD43299 (**Extended data figure 8**).

## SUPPLEMENTARY TABLES

**Note: Supplementary tables can be found in “Supplementary_tables_R0_05.01.xlsx”.**

**Supplementary table 1 | Overview of study cohort and histologies isolated for DNA and mRNA sequencing.**

**Supplementary table 2 | Overview of GCT whole genome sequencing data.**

**Supplementary table 3 | List of annotated substitutions and indels called across all GCT samples.**

**Supplementary table 4 | Copy number calls for all GCT samples that were estimated to have 5 or more reads per chromosome copy.**

**Supplementary table 5 | List of structural variants called across all GCT samples.**

**Supplementary table 6 | List of retrotransposition events called across all GCT samples.**

**Supplementary table 7 | Driver missense substitutions, their purity, and the copy number configuration at each locus.**

**Supplementary table 8 | Driver missense mutation detection across all GCT samples using CHASMplus.**

**Supplementary table 9 | Complete repertoire of SBS signatures extracted across all GCT samples.**

**Supplementary table 10 | Mutation clusters used to reconstruct the phylogenies shown in Extended data figure 6.**

**Supplementary table 11 | Proportion of invasive tumour substitutions found to lie in the trunk vs the overall burden.**

**Supplementary table 12 | Estimated pre-duplication substitution burden.**

**Supplementary table 13 | Estimation of WGD in mutation time per microbiopsy.**

**Supplementary table 14 | Overview of GCT mRNA sequencing data.**

**Supplementary table 15 | Differential gene expression between embryonal carcinoma and other GCT histologies microdissected, ranked by p value.**

**Supplementary table 16 | Gene expression enrichment by GCT tissue, binned per cytoband, relative to healthy adult seminiferous tubules.**

**Supplementary table 17 | Significantly overexpressed 12p genes, ranked by the number of invasive GCT tissues they are overexpressed in relative to normal testis.**

## DATA AVAILABILITY

The raw DNA and RNA sequencing data have been deposited in the European Genome-Phenome Archive (EGA) with the accession code EGAS00001003862.

## CODE AVAILABILITY

The R scripts used to run the bespoke filtering and analyses detailed in this study can be found here: https://github.com/trwo/GCT_diversification.

## ACKNOWLEDGEMENTS

We would like to thank Maxime Tarabichi for reviewing our copy number timing analyses, as well as Peter van Loo for providing valuable advice on our comparisons against TCGA and PCAWG reference data.

We thank the CCLG Tissue Bank, the CCLG centres and the ECMC Paediatric Network for the collection and provision of tissue samples (project numbers 2002 BS 03, 2016 BS 05 and 2020 BS 02). The CCLG Tissue Bank is funded by Cancer Research UK and CCLG. Some tissues were acquired from the Human Research Tissue Bank at Cambridge University Hospitals NHS Foundation Trust, whom we must additionally thank for their provision of histological services. The Human Research Tissue Bank is supported by the NIHR Cambridge Biomedical Research Centre. Other research samples were obtained from the Manchester Cancer Research Centre (MCRC) Biobank, UK. The role of the MCRC Biobank is to distribute samples and therefore, cannot endorse studies performed or the interpretation of results. Collection of tissues by R.H. was supported by Cancer Research UK ECMC (C9380/A25138) and the Newcastle Biobank. The CUH adult blood sampling had infrastructure support from the Urological Malignancies Programme which is part of the CRUK Cambridge Centre, funded by Cancer Research UK Major Centre Award C9685/A25117, and supported by the NIHR Cambridge Biomedical Research Centre.

Lastly, we are grateful to the patients who kindly provided the tissue and blood samples that made this study possible.

## CONTRIBUTIONS

S.B., R.R. and T.R.W.O. designed the experiment. T.R.W.O. and A.C. performed the laser capture microdissection. A.Y.W. and N.C. provided GCT histology expertise. Y.H. assisted with tissue processing and sectioning, and immunohistochemical staining. T.R.W.O. generated the final lists of substitutions, indels, copy number changes and structural variants with assistance from R.S., T.H.H.C., D.C.W. and M.S. H.J. performed the retrotransposition analysis. T.R.W.O and R.S. performed SBS signature extraction and analysis with assistance from T.H.H.C, T.B. M.D.C.N. and D.L. Copy number timing analyses were done by T.R.W.O., assisted by S.C.D. I.C. provided expertise on calling chromothripsis-like rearrangements. Phylogenetic analyses were performed by T.R.W.O. with contributions from D.C.W. and N.A.P. L.C. and R.R. designed the low-input RNA sequencing pipeline. L.C. and L.D. constructed the mRNA libraries for sequencing. RNA data processing and analysis was undertaken by T.R.W.O with guidance from M.D.Y. and R.R. M.J.M., P.J.C. and N.C. contributed to study discussions. R.H. collected study samples. S.B., T.R.W.O. and R.R. co-wrote the manuscript. S.B. and R.R. co-directed the study.

---

**Extended data figure 1.**
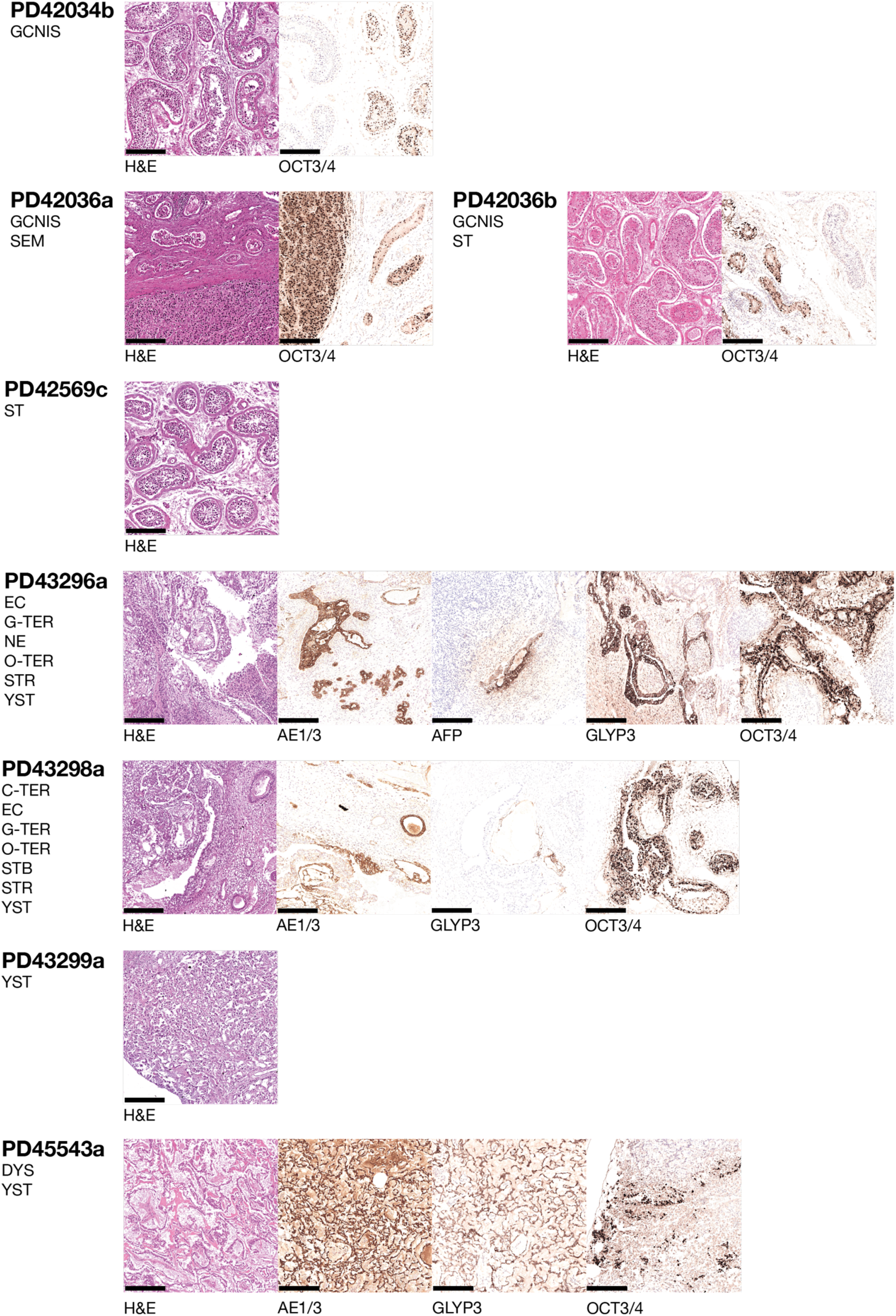
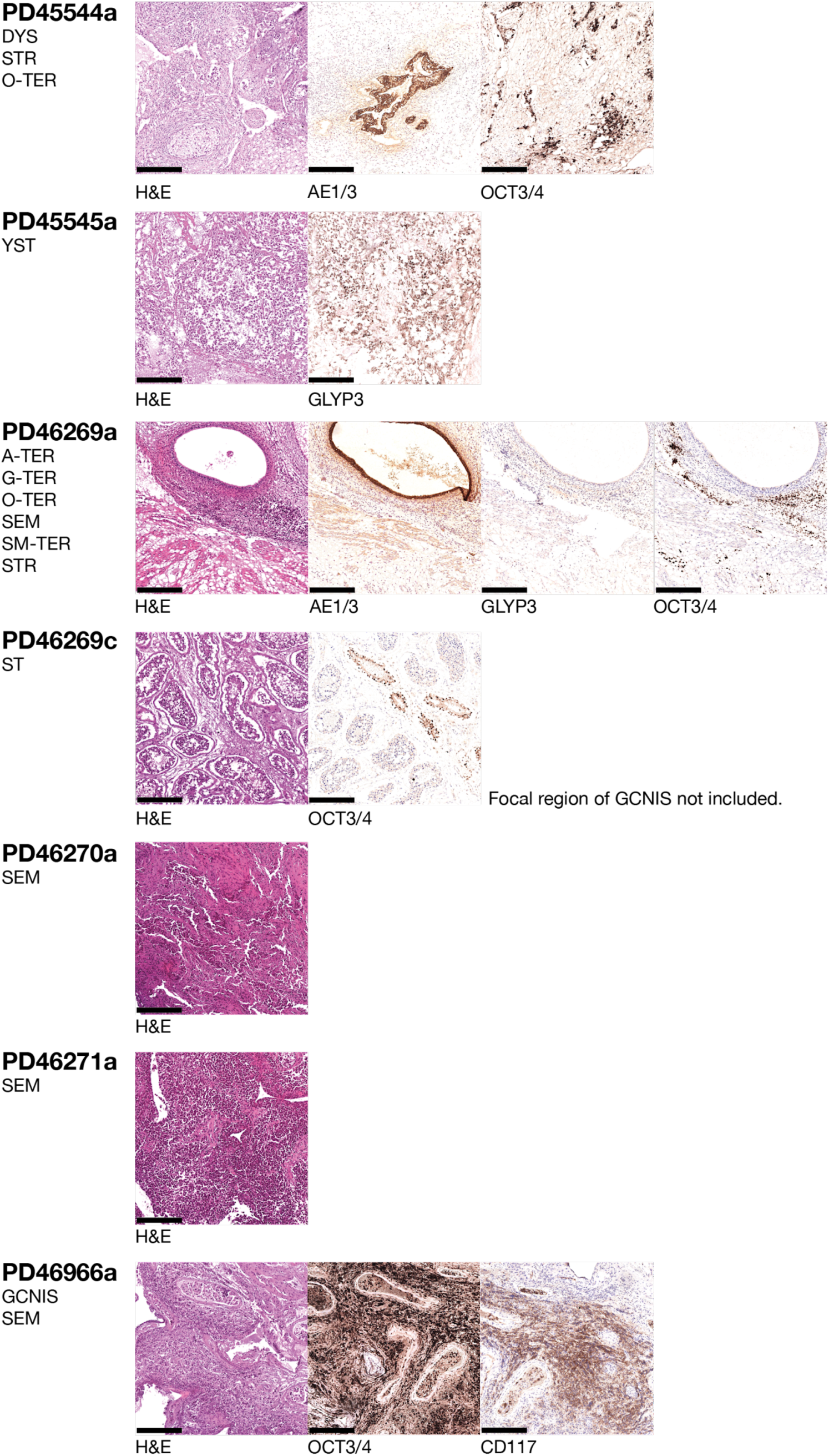
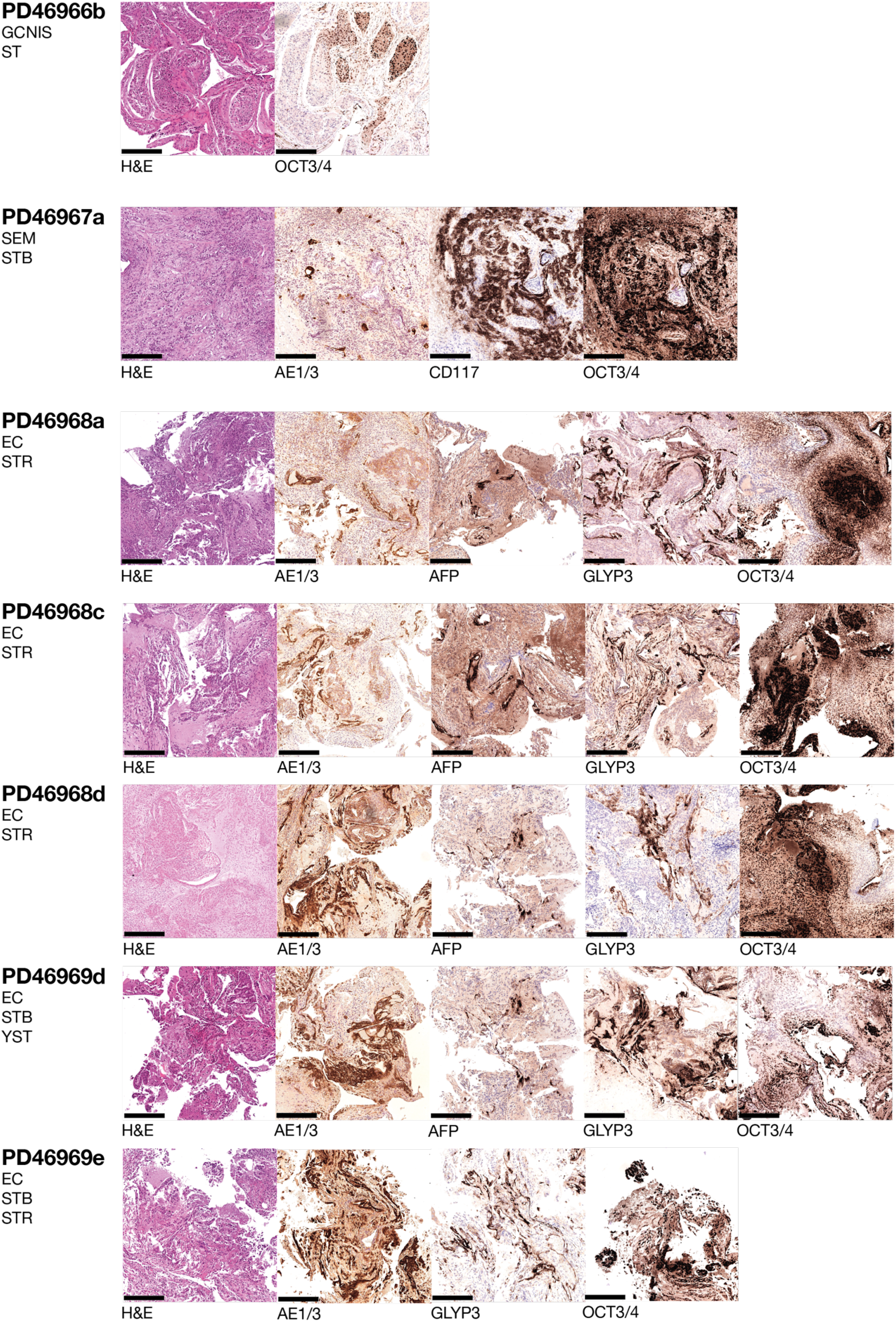
Histological images from each biopsy included in the microdissection cohort. Microphotographs taken from a reference H&E slide and any immunohistochemistry that informed histological categorisation. Underneath each case ID is a list of the histologies successfully isolated from it. For some cases, DNA and mRNA libraries were not successfully generated from all components found on the reference slides. A-TER, adipose teratoma; C-TER, cartilage teratoma; DYS, dysgerminoma; EC, embryonal carcinoma; GCNIS, germ cell neoplasia *in situ*; G-TER, mature glandular teratoma, NE, neuroepithelium; O-TER, other epithelial teratoma; SM-TER, smooth muscle teratoma; SEM, seminoma; STB, syncytiotrophoblasts; STR, malignant stroma; ST, healthy seminiferous tubules; YST, yolk sac tumour. Scale bars denote 250 microns.

**Extended data figure 2.**
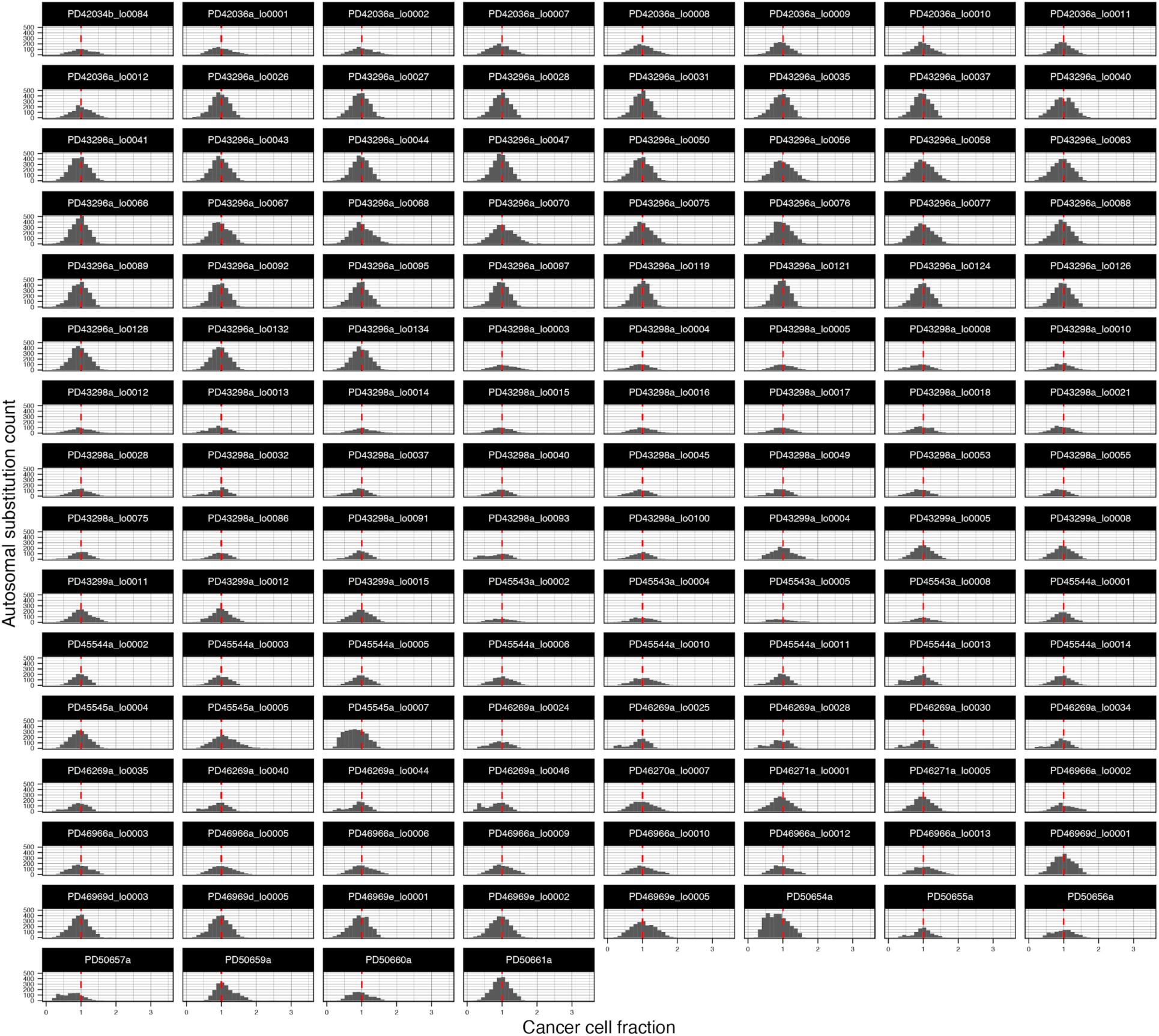
Histograms of the cancer cell fraction for every SNV lying on the autosomal genome in each GCT. Only samples with an estimated minimum of five reads per chromosome copy were included, reflecting the threshold used to have sufficient confidence to include the associated copy number data in downstream analyses.

**Extended data figure 3.**
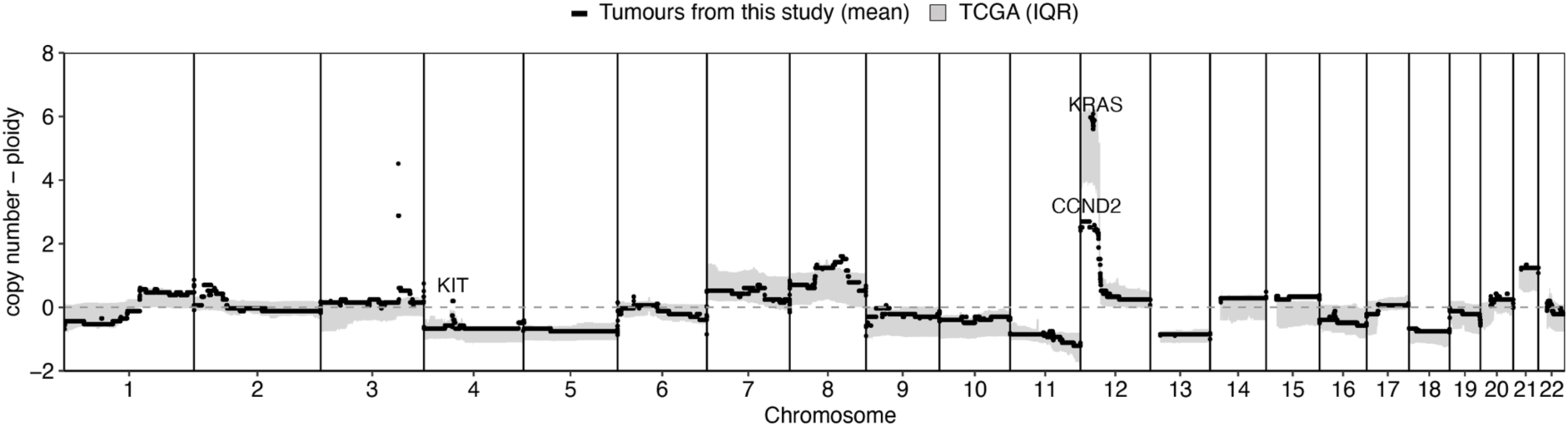
Aggregated copy number across the microdissection cohort compared to TCGA reference data [6]. The study cohort average excludes the single prepubertal case (PD43299) as none such cases were present in the TCGA dataset.

**Extended data figure 4.**
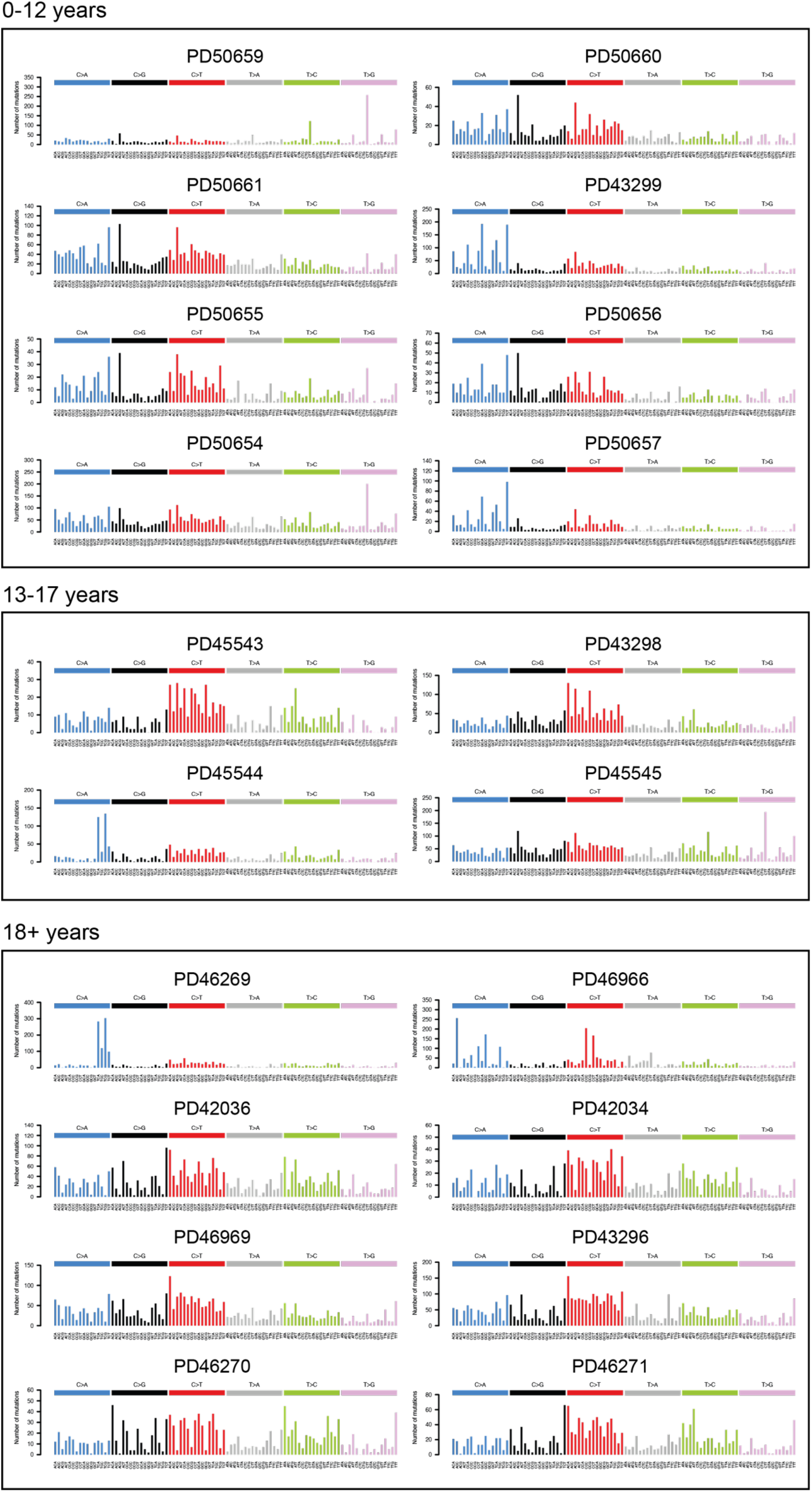
Trinucleotide context plots of all unique substitutions per GCT. Each GCT is arranged by age.

**Extended data figure 5.**
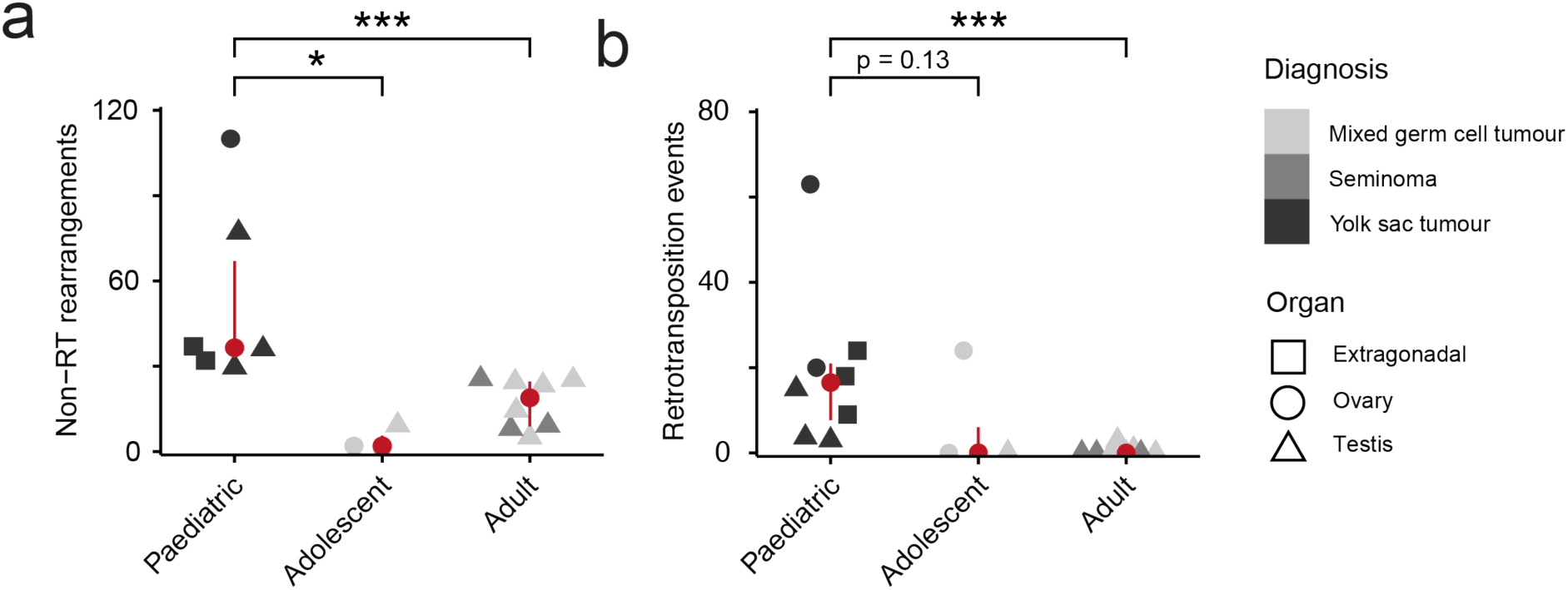
Structural variant burden per GCT. (**a**) Burden of all structural variants except retrotransposition (RT) events. Cases which have undergone a chromothripsis-like event are excluded. Cases are binned by age; “Paediatric” (0-12 years), “Adolescent” (13-17 years) and “Adult” (18+ years). P values were acquired using the Wilcoxon rank sum test (paediatric vs adult; p <0.001). (**b**) Retrotransposition events called per tumour, binned as per (**a**). P values were acquired using the Wilcoxon rank sum test (paediatric vs adult; p <0.001).

**Extended data figure 6.**
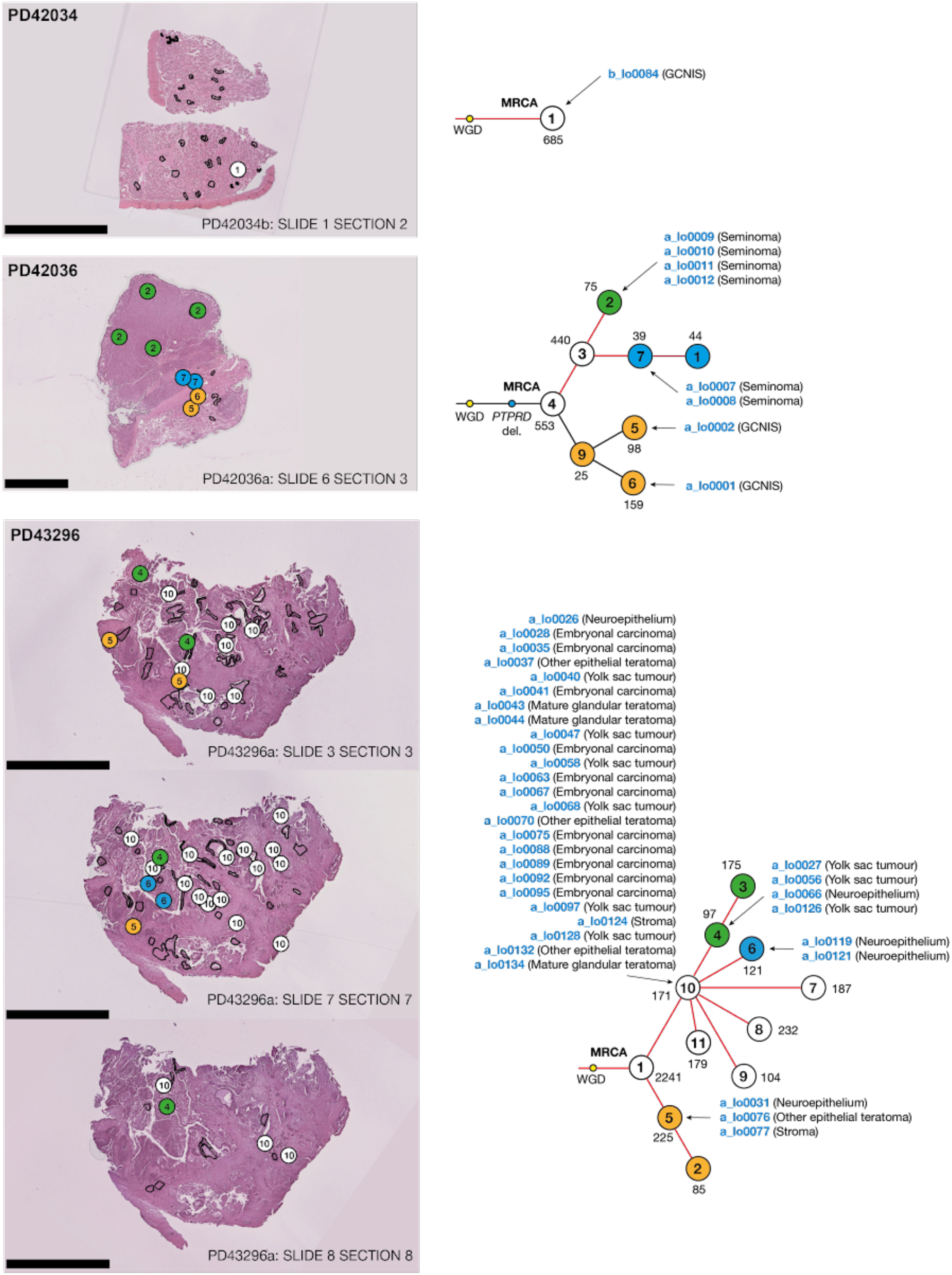
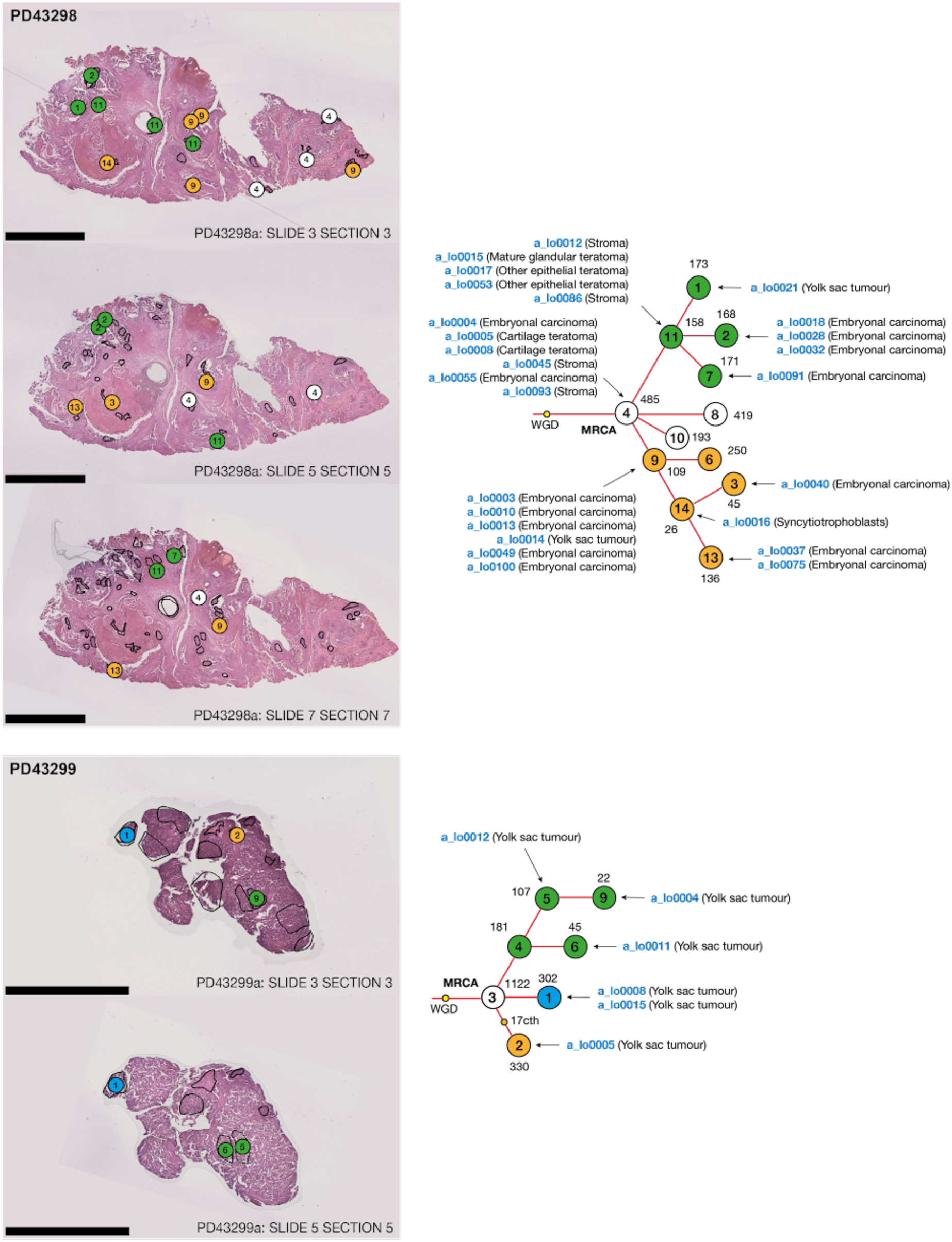
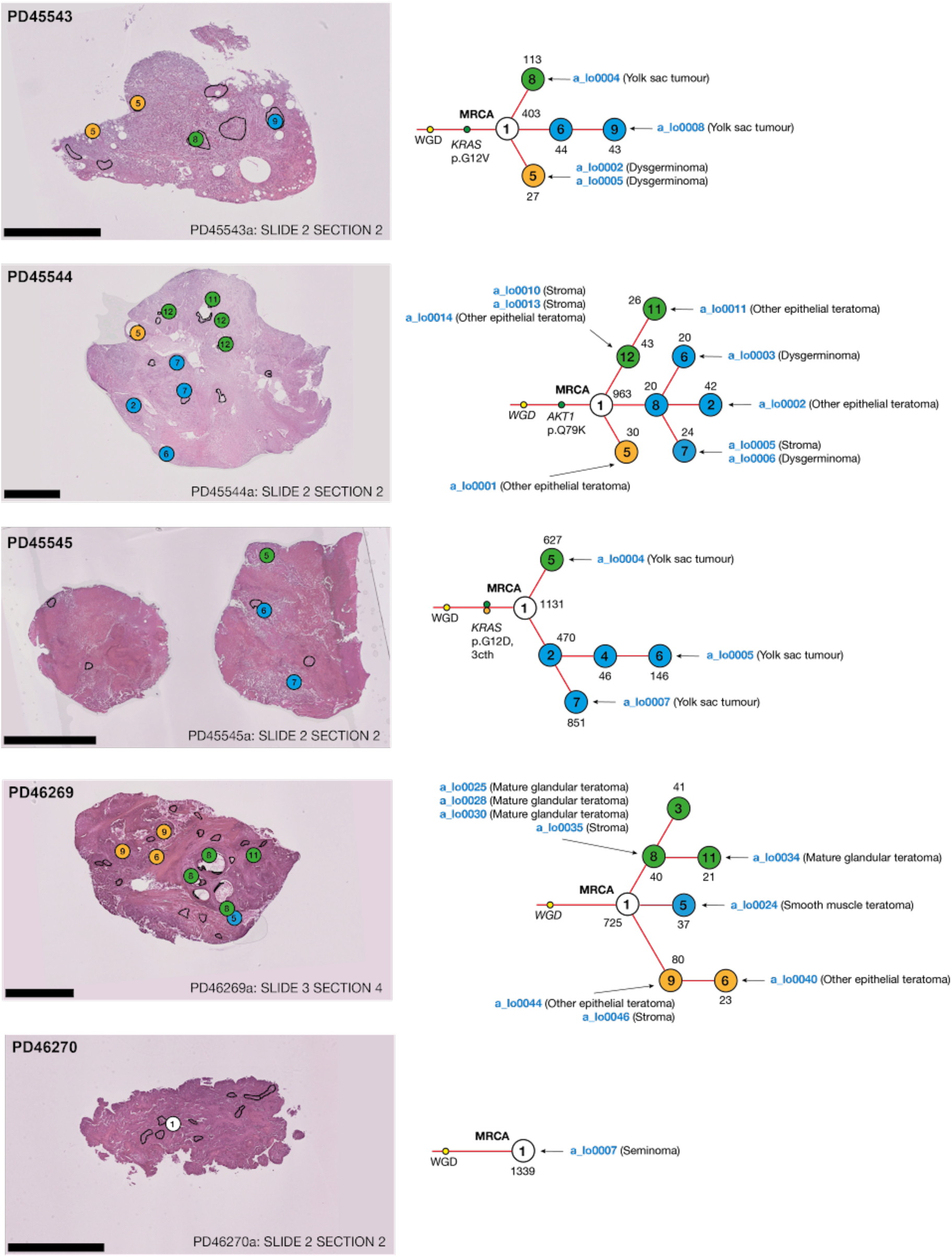
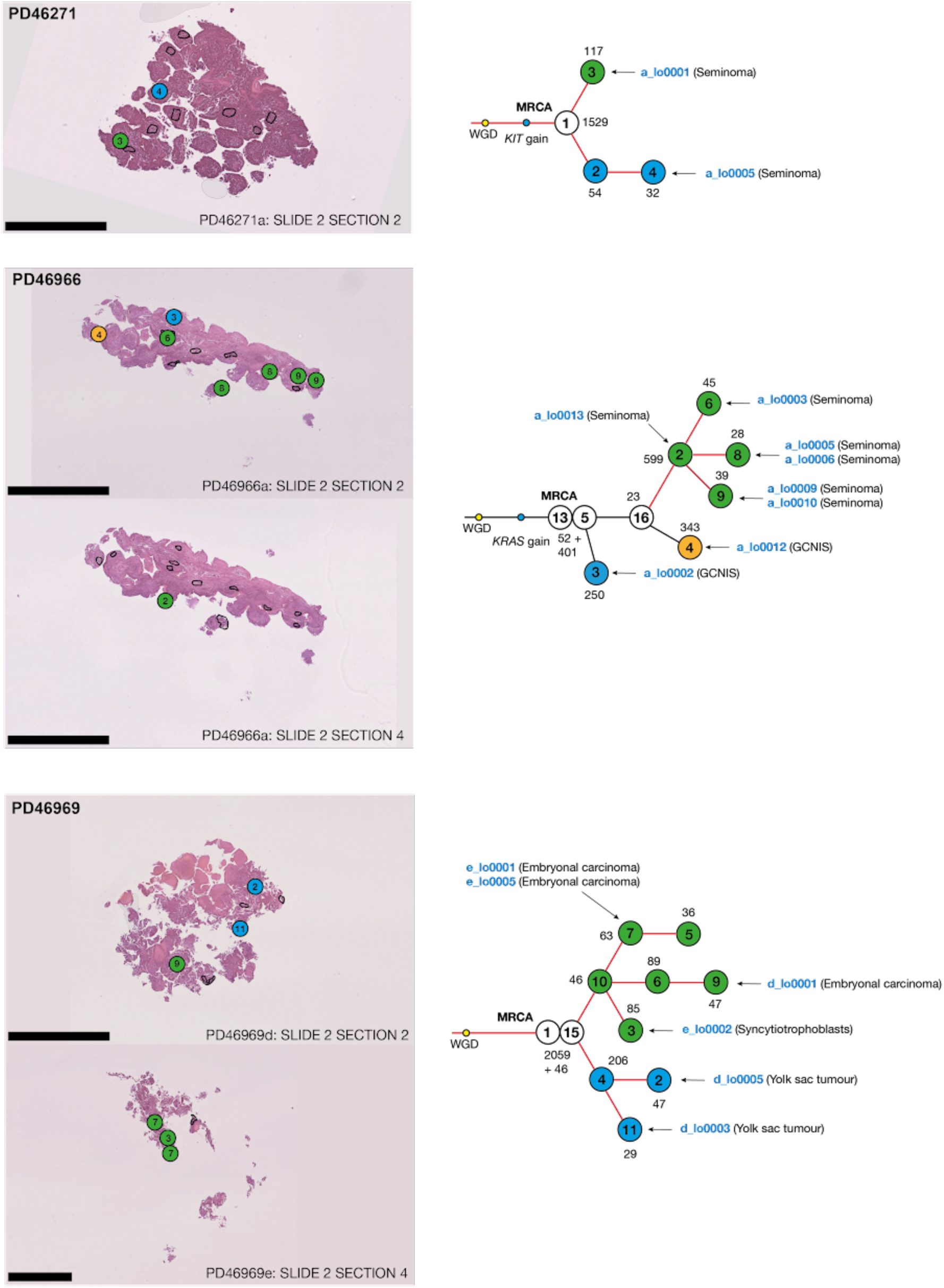
GCT phylogenies mapped back to histological sections. Each circle represents a mutation cluster. The number within the circle is the arbitrary cluster number provided by the phylogeny algorithm. The number adjacent to each circle is the number of autosomal SNVs supporting it. Black lines between clusters denote an *in situ* relationship whereas red lines represent linked ancestry following invasion. Each eligible microbiopsy is attributed to the mutation cluster that is the dominant clone (>0.5 cancer cell fraction) found within it. The scale bar for PD42034 indicates 10mm, for PD45545 it is 5mm and for the remainder it is 2.5mm.

**Extended data figure 7.**
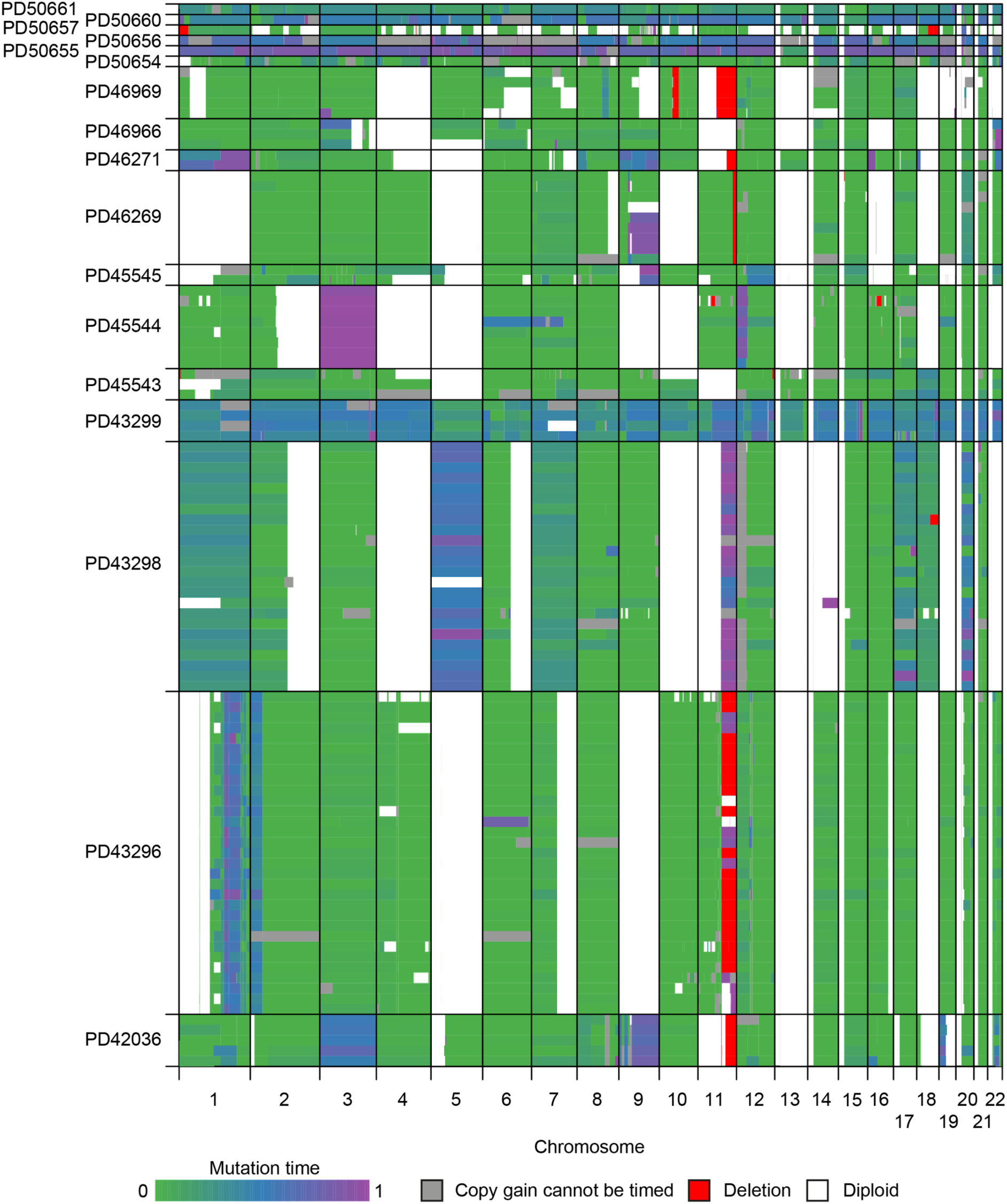
Overview of the mutationtimeR estimates for each copy gain called per GCT sample. Complex, high copy gains and deletions cannot be timed and thus are coloured grey and red respectively. If a segment is gained multiple times and timing estimates are still available, the segment is coloured according to the first gain.

**Extended data figure 8.**
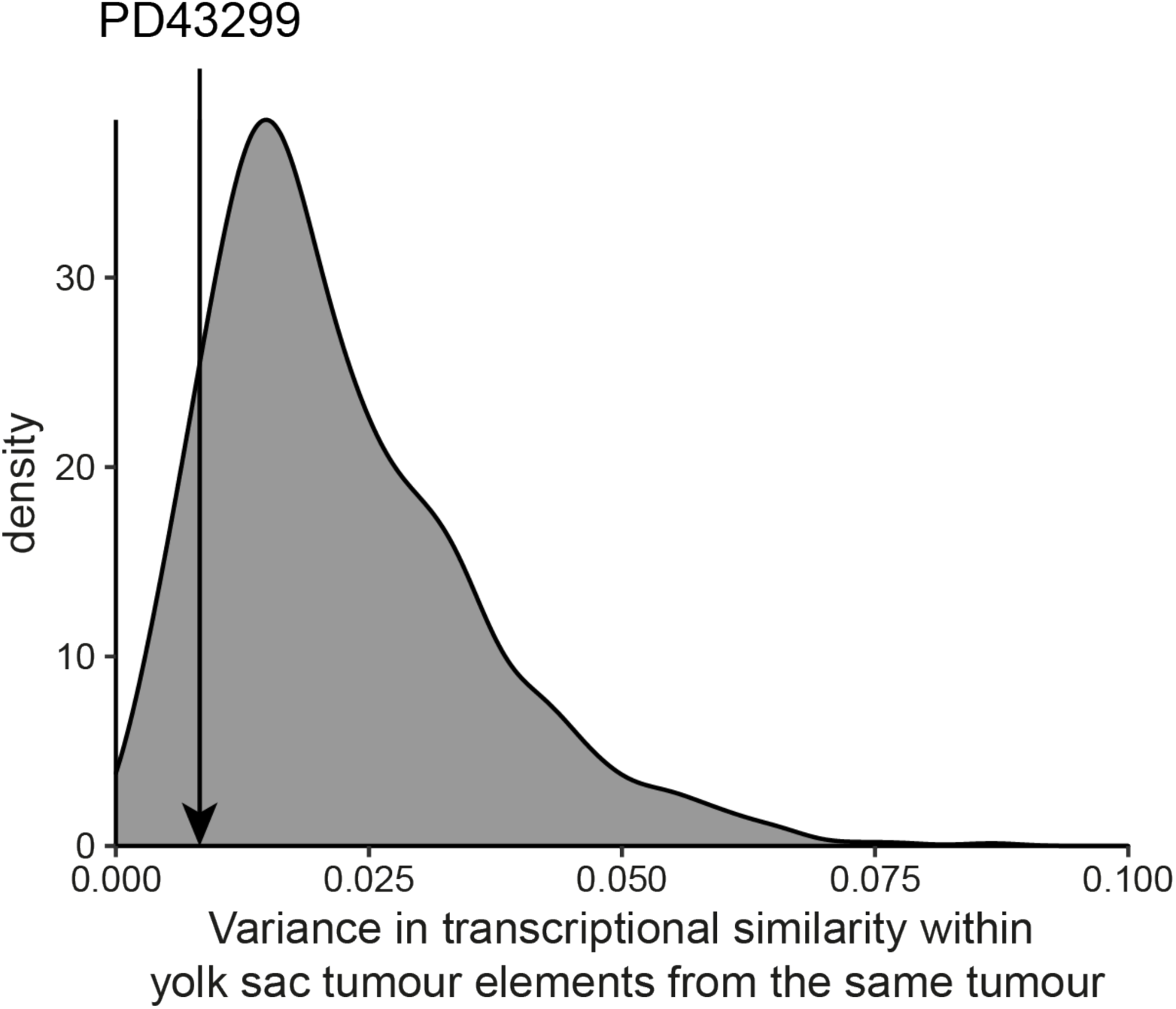
Intra-tumoral yolk sac tumour transcriptional heterogeneity. Pairwise Pearson correlation of the transcriptome of yolk sac tumour elements within each tumour was calculated before the variance of 1,000 random subsets was calculated to achieve the distribution illustrated here. This provides an estimate of the transcriptional variability across the yolk sac tumour within a given tumour. PD43299 did not show a greater transcriptional variability compared to the cohort overall, despite subclonal chromothripsis of chromosome 17.

**Extended data figure 9.**
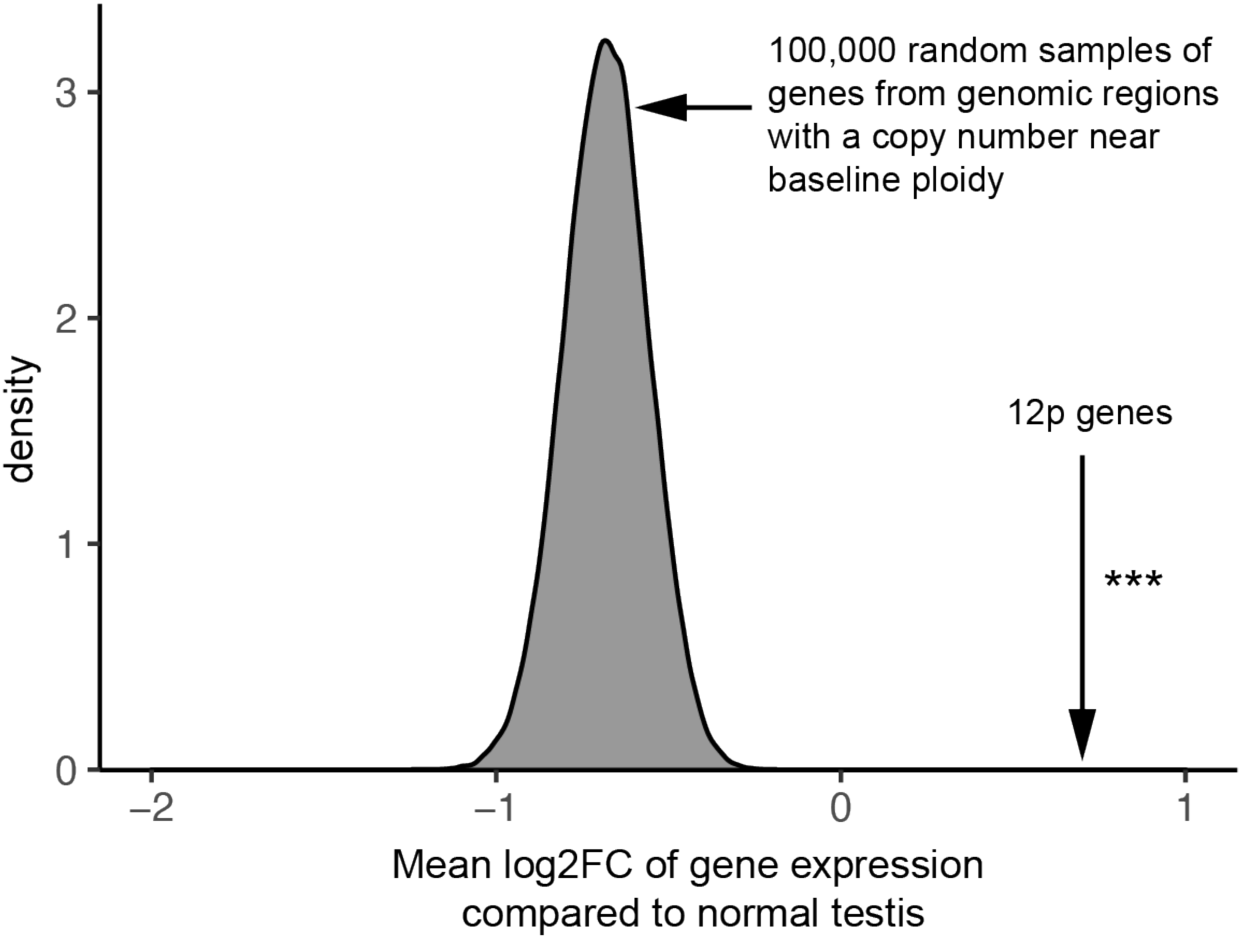
The distribution of the mean log2 fold change in expression compared to normal testis tissues across 100,000 random subsets of genes from GCT genomic regions near baseline ploidy vs 12p genes. No random samples achieved a mean log2 fold change compared to healthy testis higher than that achieved across 12p (p < 10^-5^).

**Extended data figure 10.**
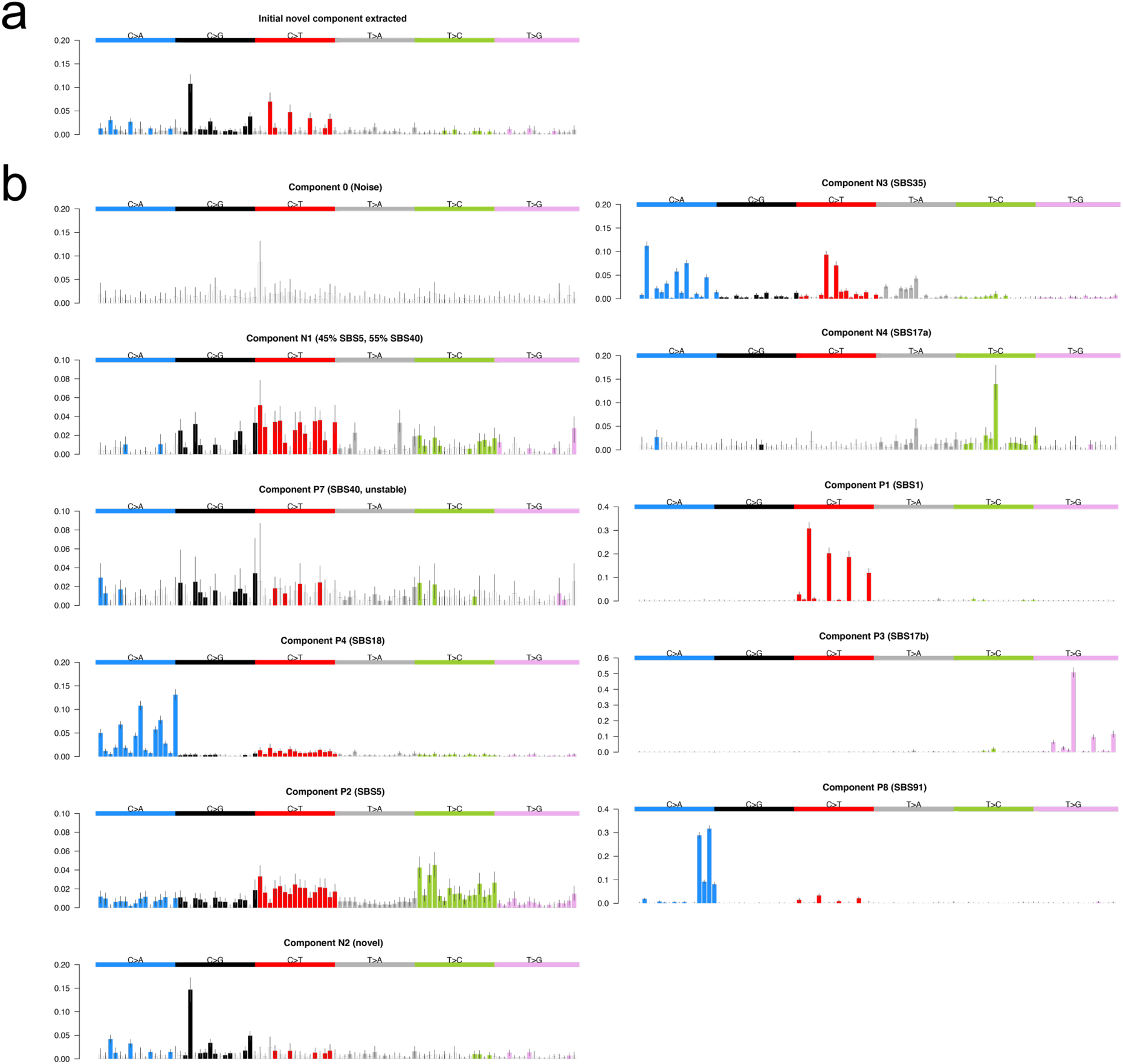
HDP signature extraction components. (**a**) The initial novel component extracted using the entire list of COSMIC version 3.2 signatures. The C>T peaks indicated that it had been extracted with an element of SBS1. (**b**) The final list of components extracted and the signatures they corresponded to. The whiskers on each bar represent the 95% credible interval.

**Extended data figure 11.**
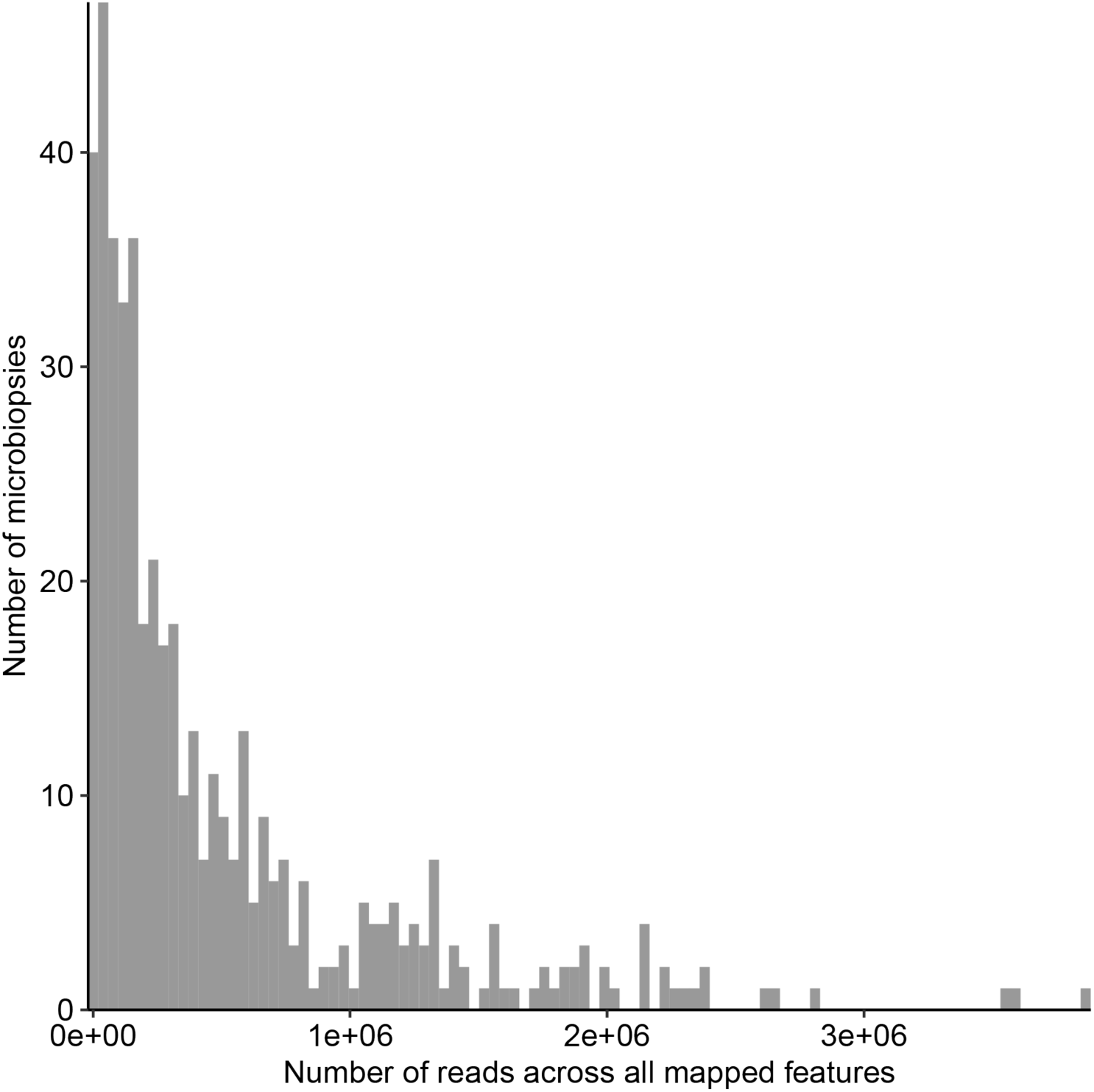
Distribution of the pre-QC total read depth for all GCT transcriptomes. The library size for most microbiopsies are on a scale somewhere between single cell and standard bulk RNA sequencing.

**Extended data figure 12.**
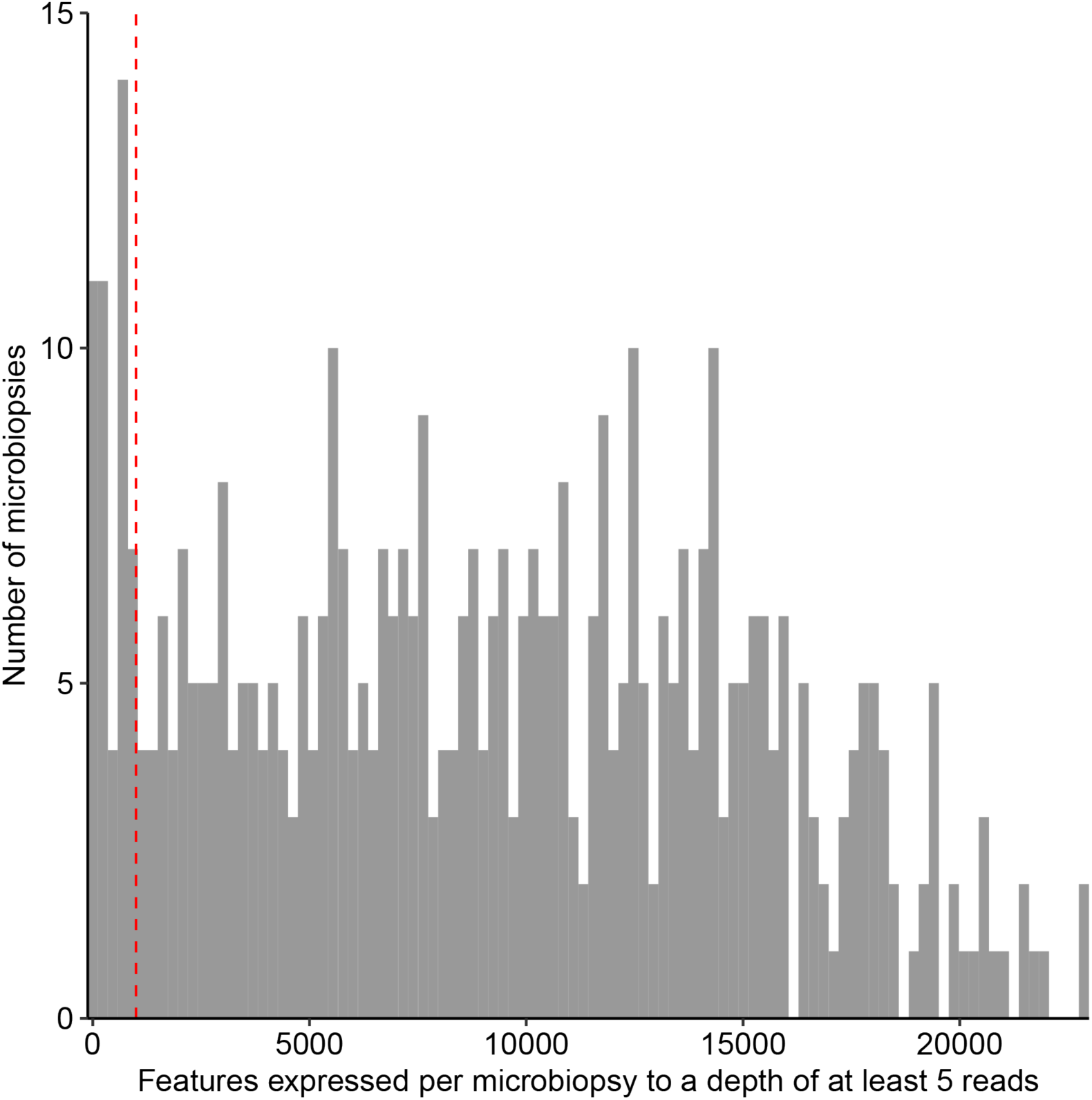
Distribution of the number of genes expressed to a depth of at least 5 reads for all pre-filtered transcriptomes. The dashed red line is drawn at 1,000 genes, the threshold applied to remove low quality samples.

**Extended data figure 13.**
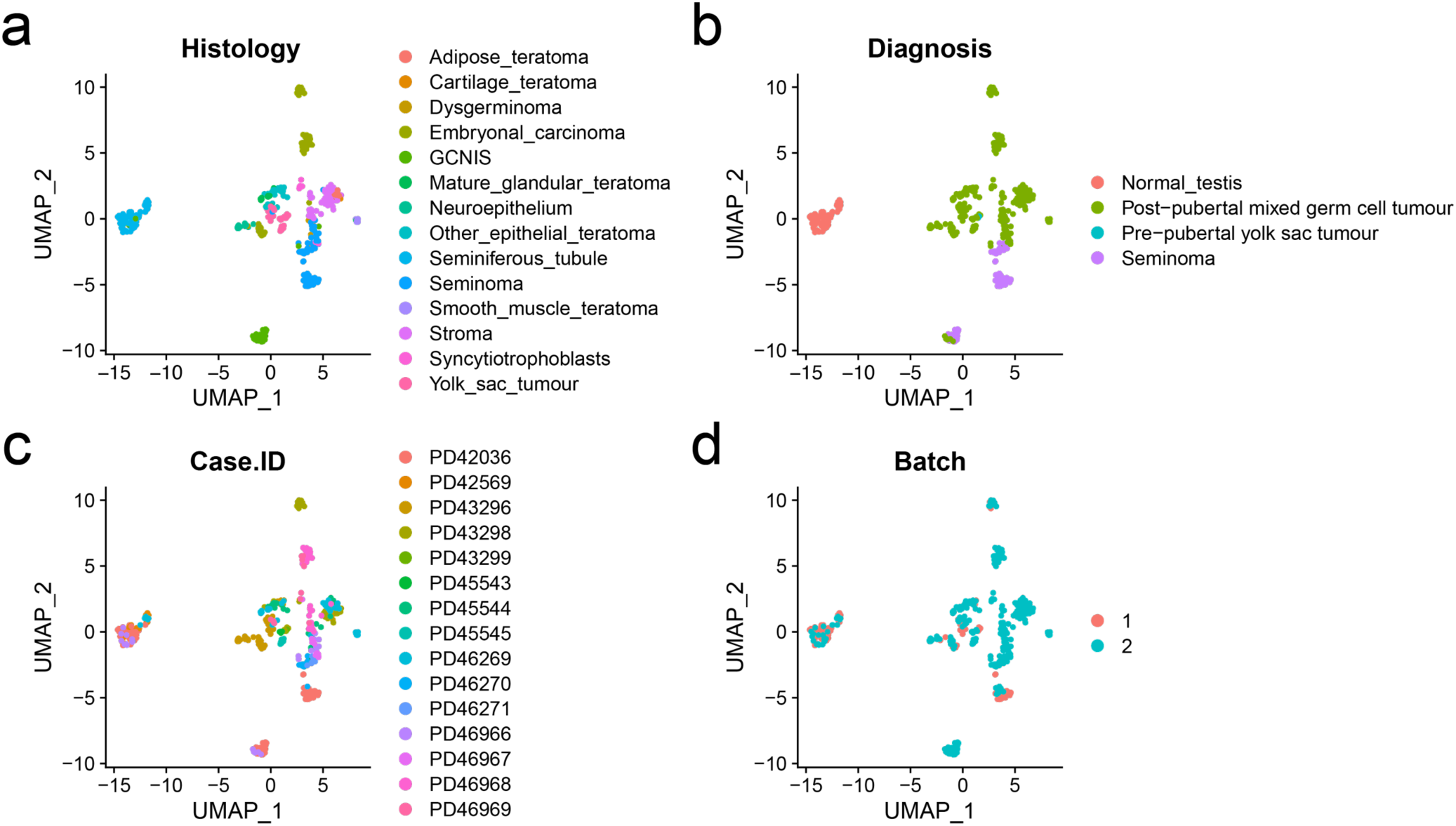
UMAP clustering of RNA microbiopsy data. Coloured by histology (**a**), tumour diagnosis (**b**), case ID (**c**) and batch the samples were submitted in (**d**). The first 30 dimensions were chosen during the creation of the UMAP.

**Extended data figure 14.**
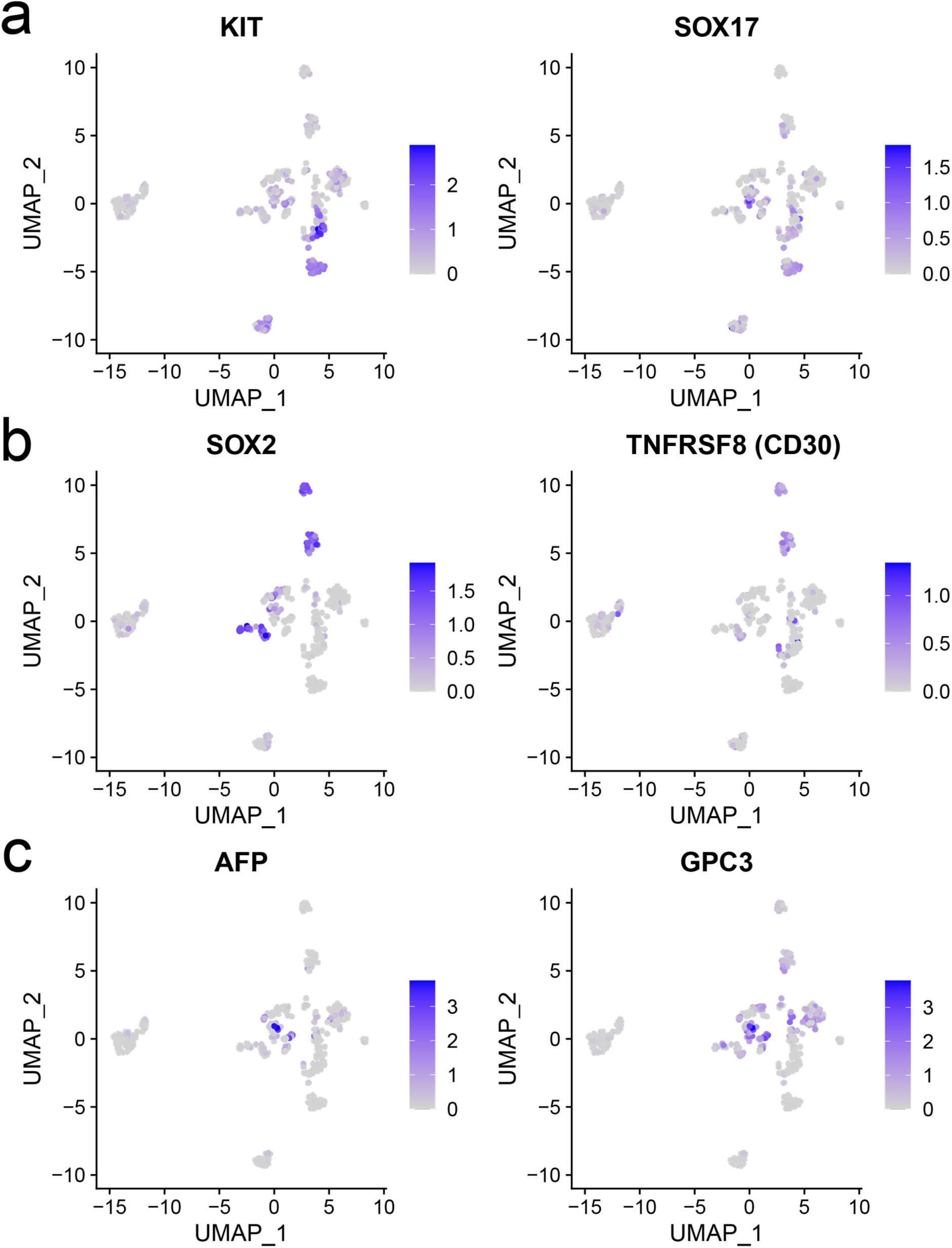
Normalised expression of marker genes that define GCT histologies. Marker genes were chosen for seminoma (**a**), embryonal carcinoma (**b**) and yolk sac tumour (**c**); the three histologies best characterised from previous studies [72–75].

## REFERENCES

1. Trabert, B., Chen, J., Devesa, S. S., Bray, F. & McGlynn, K. A. International patterns and trends in testicular cancer incidence, overall and by histologic subtype, 1973-2007. Andrology 3, 4–12 (2015).

2. Williamson, S. R. et al. The World Health Organization 2016 classification of testicular germ cell tumours: a review and update from the International Society of Urological Pathology Testis Consultation Panel. Histopathology 70, 335–346 (2017).

3. International Germ Cell Cancer Collaborative Group. International Germ Cell Consensus Classification: a prognostic factor-based staging system for metastatic germ cell cancers. International Germ Cell Cancer Collaborative Group. J. Clin. Oncol. 15, 594–603 (1997).

4. Murray, M.J. et al. Treatment and outcomes of UK and German patients with relapsed intracranial germ cell tumors following uniform first-line therapy. Int. J. Cancer 141, 621–635 (2017).

5. Karlsson, J. et al. Four evolutionary trajectories underlie genetic intratumoral variation in childhood cancer. Nat. Genet. 50, 944–950 (2018).

6. Shen, H. et al. Integrated Molecular Characterization of Testicular Germ Cell Tumors. Cell Rep. 23, 3392–3406 (2018).

7. Taylor-Weiner, A. et al. Genomic evolution and chemoresistance in germ-cell tumours. Nature 540, 114–118 (2016).

8. Litchfield, K. et al. Whole-exome sequencing reveals the mutational spectrum of testicular germ cell tumours. Nat. Commun. 6, 5973 (2015).

9. Dorssers, L. C. J. et al. Molecular heterogeneity and early metastatic clone selection in testicular germ cell cancer development. Br. J. Cancer 120, 444–452 (2019).

10. Loveday, C. et al. Genomic landscape of platinum resistant and sensitive testicular cancers. Nat. Commun. 11, 2189 (2020).

11. Van Nieuwenhuysen, E. et al. The genetic landscape of 87 ovarian germ cell tumors. *Gynaecol*. Oncol. 151, 61–68 (2018).

12. Atkin, N. B. & Baker, M.C. Specific chromosome change, i(12p), in testicular tumours? Lancet 11, 1349 (1982).

13. Coorens, T.H.H. et al. Somatic mutations reveal widespread mosaicism and mutagenesis in human placentas. Nature 592, 80–85 (2021).

14. Gröbner, S. N. et al. The landscape of genomic alterations across childhood cancers. Nature 555, 321–327 (2018).

15. Alexandrov, L. B. et al. The repertoire of mutational signatures in human cancer. Nature 578, 94–101 (2020).

16. Gerstung, M. et al. The evolutionary history of 2,658 cancers. Nature 578, 122–128 (2020).

17. Rahbari, R. et al. Timing, rates and spectra of human germline mutation. Nat. Genet. 48, 126–133 (2016).

18. Behjati, S. et al. Recurrent mutation of IGF signalling genes in distinct patterns of genomic rearrangement in osteosarcoma, Nat. Commun. 8, 15936 (2017).

19. Sohni, A. et al. The Neonatal and Adult Human Testis Defined at the Single-Cell Level. Cell rep. 26, 1501–1517 (2019).

20. Zhong, S. et al. A single-cell RNA-seq survey of the developmental landscape of the human prefrontal cortex. Nature 555, 524–528 (2018).

21. Li, M. et al. Integrative functional genomic analysis of human brain development and neuropsychiatric risks. Science 362, eaat7615 (2018).

22. Cao, J. et al. A human cell atlas of fetal gene expression. Science 370, eaba7721 (2020).

23. Litviňuková, M. et al. Cells of the adult human heart. Nature 588, 466–472 (2020).

24. White, V. et al. IGF2 stimulates fetal growth in a sex- and organ-dependent manner. Pediatr. Res. 83, 183–189 (2018).

25. McBurney, M., Jonas-Villeneuve, E. M. V., Edwards, M. K. S. & Anderson, P. J. Control of muscle and neuronal differentiation in a cultured embryonal carcinoma cell line. Nature 299, 165–167 (1982).

26. Skotheim R.I. et al. Differentiation of human embryonal carcinomas in vitro and in vivo reveals expression profiles relevant to normal development. Cancer Res. 65, 5588–5598 (2005).

27. Xia, X. & Su-Chun, Z. Differentiation of neuroepithelia from human embryonic stem cells. Methods Mol. Biol. 549, 51–8 (2009).

28. Dee, C. T. et al. Sox3 regulates both neural fate and differentiation in the zebrafish ectoderm. Dev. biol. 320, 289–301 (2008).

29. Wang, Z., Wang, D. Z., Teg Pipes, G. C. & Olson, E. N. Myocardin is a master regulator of smooth muscle gene expression. Proc. Natl Acad. Sci. U. S. A. 100, 7129–34 (2003).

30. Cordes, K. et al. miR-145 and miR-143 regulate smooth muscle cell fate and plasticity. Nature 460, 705–710 (2009).

31. Looijenga, L. H., Gillis, A. J., Stoop, H. J., Hersmus, R. & Oosterhuis, J. W. Chromosomes and expression in human testicular germ-cell tumors: insight into their cell of origin and pathogenesis. Ann N Y Acad Sci. 2007 1120, 187–214 (2007).

32. Korkola, J. E. et al. Down-regulation of stem cell genes, including those in a 200-kb gene cluster at 12p13.31, is associated with in vivo differentiation of human male germ cell tumors. Cancer Res. 66, 820–7 (2006).

33. Shao, X. et al. Copy number variation is highly correlated with differential gene expression: a pan-cancer study. BMC Med Genet. 20, 175 (2019).

34. Martic, G. et al. Parathymosin affects the binding of linker histone H1 to nucleosomes and remodels chromatin structure. J. Biol. Chem. 280, 16143–16150 (2005).

35. Huang, M. et al. Use of all trans retinoic acid in the treatment of acute promyelocytic leukaemia. Blood 72, 567–572 (1988).

36. Degos, L. et al. All-trans retinoic acid as a differentiating agent in the treatment of acute promyelocytic leukemia. Blood 85, 2643–2653 (1995).

37. Villablanca, J. G. et al. Phase I trial of 13-cis-retinoic acid in children with neuroblastoma following bone marrow transplantation. J. Clin. Oncol. 13, 894–901 (1995).

38. Matthay, K. K., et al. Children’s Cancer Group. Treatment of high-risk neuroblastoma with intensive chemotherapy, radiotherapy, autologous bone marrow transplantation, and 13-cis- retinoic acid. N. Engl. J. Med. 341, 1165–1173 (1999).

39. Hayashi, S. et al. Primary non-gestational pure choriocarcinoma arising in the ovary: A case report and literature review. Oncol Lett. 9, 2109–2111 (2015).

40. Oosterhuis, J. W. & Looijenga, L. H. J. Human germ cell tumours from a developmental perspective. Nat Rev Cancer 19, 522–537 (2019).

41. Schneider, D. T. et al. Epidemiologic analysis of 1,442 children and adolescents registered in the German germ cell tumor protocols. Pediatr Blood Cancer 42, 169–175 (2004).

42. Calaminus, G., et al. Age-Dependent Presentation and Clinical Course of 1465 Patients Aged 0 to Less than 18 Years with Ovarian or Testicular Germ Cell Tumors; Data of the MAKEI 96 Protocol Revisited in the Light of Prenatal Germ Cell Biology. Cancers (Basel) 12, 611 (2020).

43. Palmer, R. D. et al. Malignant germ cell tumours of childhood: new associations of genomic imbalance. Br. J. Cancer. 96, 667–676 (2007).

44. Palmer, R. D. et al. Pediatric malignant germ cell tumors show characteristic transcriptome profiles. Cancer Res. 68, 4239–47 (2008).

45. Shaikh, F. et al. Paediatric extracranial germ-cell tumours. Lancet Oncol. 17, e149–e162 (2016).

## METHODS REFERENCES

46. Ellis, P. et al. Reliable detection of somatic mutations in solid tissues by laser-capture microdissection and low-input DNA sequencing. Nat Protoc 16, 841–871 (2021).

47. Li H., Durbin R., Fast and accurate short read alignment with Burrows–Wheeler transform. Bioinformatics, 25, 1754–1760, (2009).

48. Picelli, S. et al. Full-length RNA-seq from single cells using Smart-seq2. Nat Protoc 9, 171–181 (2014).

49. Dobin, A. et al. STAR: ultrafast universal RNA-seq aligner. Bioinformatics 29, 15–21 (2013).

50. Bergmann, E. A., Chen, B. J., Arora, K., Vacic, V. & Zody, M. C. Conpair: concordance and contamination estimator for matched tumor-normal pairs. Bioinformatics 32, 3196–3198 (2016).

51. Pedersen, B. S. & Quinlan, A. R. Mosdepth: quick coverage calculation for genomes and exomes. Bioinformatics 34, 867–868 (2018).

52. Jones, D. et al. cgpCaVEManWrapper: simple execution of CaVEMan in order to detect somatic single nucleotide variants in NGS data. Curr. protoc. Bioinformatics 56, 15.10.1–15.10.18 (2016).

53. Coorens, T. H. H. et al. Embryonal precursors of Wilms tumor. Science 366, 1247–1251 (2019).

54. Moore, L. et al. The mutational landscape of human somatic and germline cells. Nature 597, 381–386 (2021).

55. Coorens, T. H. H. et al. Extensive phylogenies of human development inferred from somatic mutations. Nature 597, 387–392 (2021).

56. Buels, R. et al. JBrowse: a dynamic web platform for genome visualization and analysis. Genome biology, 17(1), 66, (2016).

57. Ye, K., Schulz, M.H., Long, Q., Apweiler, R. & Ning, Z. Pindel: a pattern growth approach to detect break points of large deletions and medium sized insertions from paired-end short reads. Bioinformatics 25, 2865–2871 (2009).

58. Nik-Zainal, S. et al. The life history of 21 breast cancers. Cell 149, 994–1007 (2012).

59. Moore, L. et al. The mutational landscape of normal human endometrial epithelium. Nature 580, 640–646 (2020).

60. Rodriguez-Martin, B. et al. Pan-cancer analysis of whole genomes identifies driver rearrangements promoted by LINE-1 retrotransposition. Nat. Genet. 52, 306–319 (2020).

61. Teh, Y. W., Jordan, M. I., Beal, M. J. & Blei, D. M. Hierarchical Dirichlet Processes. J. Am. Stat. Assoc. 101, 1566–1581 (2006).

62. Robinson, P.S. et al. Elevated somatic mutation burdens in normal human cells due to defective DNA polymerases. bioRxiv, (2020).

63. Martincorena, I. et al. Universal Patterns of Selection in Cancer and Somatic Tissues. Cell 171, 1029–1041.e21 (2017).

64. Tokheim, C. & Karchin R. CHASMplus Reveals the Scope of Somatic Missense Mutations Driving Human Cancers. Cell Syst. 9, 9–23.e8 (2019).

65. Cortés-Ciriano, I. et al. Comprehensive analysis of chromothripsis in 2,658 human cancers using whole-genome sequencing. Nat Genet 52, 331–341 (2020).

66. Rabbie, R. et al. Multi-site clonality analysis uncovers pervasive heterogeneity across melanoma metastases. Nat Commun 11, 4306 (2020).

67. Behjati, S. et al. Genome sequencing of normal cells reveals developmental lineages and mutational processes. Nature 513, 422–425, (2014).

68. Ju, Y. S. et al. Somatic mutations reveal asymmetric cellular dynamics in the early human embryo. Nature 543, 714–718, (2017).

69. Netto, G. J. et al. Global DNA hypomethylation in intratubular germ cell neoplasia and seminoma, but not in nonseminomatous male germ cell tumors. Mod. Pathol. 21, 1337–1344 (2008).

70. Liao, Y., Smyth, G. K. & Shi, W. featureCounts: an efficient general purpose program for assigning sequence reads to genomic features. Bioinformatics. 30, 923–930 (2014).

71. Hao, Y. et al. Integrated analysis of multimodal single-cell data. Cell. 184, 3573–3587 (2021).

72. Nikolaou, M. et al. Kit expression in male germ cell tumors. Anticancer Res. 27, 1685–1688 (2007).

73. Nonaka, D. Differential expression of SOX2 and SOX17 in testicular germ cell tumors. Am J Clin Pathol. 131, 731–736 (2009).

74. Pallesen, G. & Hamilton-Dutoit, S. J. Ki-1 (CD30) antigen is regularly expressed by tumor cells of embryonal carcinoma. Am J Pathol 133, 446–450 (1988).

75. Zynger, D. L., McCallum, J. C., Luan, C., Chou, P. M., Yang, X. J. Glypican 3 has a higher sensitivity than alpha-fetoprotein for testicular and ovarian yolk sac tumour: immunohistochemical investigation with analysis of histological growth patterns. Histopathology. 56, 750–757 (2010).

76. Ritchie, M. E. et al. Limma powers differential expression analyses for RNA-sequencing and microarray studies. Nucleic Acids Res. 43, e47 (2015).

77. Subramanian, A. et al. Gene set enrichment analysis: a knowledge-based approach for interpreting genome-wide expression profiles. Proc. Natl. Acad. Sci. U. S. A. 102, 15545–15550 (2005).

